# Exploration of mScarlet for development of a red lifetime sensor for calcium imaging

**DOI:** 10.1101/2024.12.22.628354

**Authors:** Franka H. van der Linden, Theodorus W.J. Gadella, Joachim Goedhart

## Abstract

The past decades, researchers have worked on the development of genetically encoded biosensors, including over 60 genetically encoded calcium indicators (GECIs) containing a single fluorescent protein (FP). Red fluorescent GECIs provide advantages in terms of imaging depths and reduced cell toxicity. Most of GECIs respond with a fluorescence intensity change, and researchers have strived to improve the sensors in terms of brightness and fold-change. Unfortunately, fluorescence intensity is influenced by many factors other than the desired sensor response. GECIs with a fluorescence lifetime contrast overcome this drawback, but so far, no bright red GECI has been developed that shows a fluorescence lifetime contrast. We tried to tackle this challenge by using the brightest red fluorescent proteins from the mScarlet family to develop a new sensor. We did succeed in creating remarkable bright probes, but the fluorescence lifetime contrast we observed in bacterial lysates was lost in mammalian cells. Based on our results, and the success of others to develop a pH and a voltage sensor of mScarlet, we are confident that a GECI with mScarlet is feasible. To this end, we propose to continue development using a mammalian cell-based screening, instead of screening in bacterial lysates.

## Introduction

The family of single fluorescent proteins (FP) based biosensors for calcium consists currently of over 60 members ^1^. Initially, yellow FPs were used as basis for these sensors, followed quickly by green variants ^2–5^. Later, other colors were added to the palette, including the first red variants ^6,7^. The red color is superior to blue shifted colors in terms of phototoxicity and tissue penetration. However, it comes at a price: like red FPs, red calcium sensors are more difficult to detect than their green counterparts due to a (much) lower brightness.

So far, mApple is the most widely used red FP in calcium sensors (in R-GECO1^6^, R-CaMP1.07^7^, R-GECO1.2^8^, O-GECO^8^, CAR-GECO^8^, LAR-GECO^8^, R-CEPIA1er ^9^, R-CaMP2^10^, RGECO1a ^11^ and FR-CaMP^12^). Also mRuby (in the RCaMP^13^ and jRCaMP^11^ series), FusionRed (in K-GECO1^14^) and mCherry (in CH-GECO^15^) have been exploited, although the latter contains an accidental mix-up of mApple and mCherry. The past years, improved and brighter red FPs have been developed, with the mScarlet series being the brightest ^16,17^. These FPs gain their brightness due to fast maturation and a high quantum yield (QY). Potentially, using a mScarlet variant could lead to a bright calcium sensor. Others have successfully created a pH sensor ^18^ and a voltage sensor ^19^ based on mScarlet, both showing an improvement in brightness over other red versions.

In addition to brightness, the mode of read-out is also of interest. Until now, all red sensors have been optimized for an intensity contrast between absence and presence of calcium. As intensity is influenced by other factors, as cell thickness, cell movement, expression level, instrumentation and settings, this leads to difficulty when quantifying the signal. Using a fixed property of the fluorescence, like the fluorescence life-time, circumvents these issues. RCaMP1h has been successfully used for lifetime-based calcium sensing ^20^, and we showed that jRCaMP1b has the same potential ^21^. However, both these sensors are dim, especially in the calcium-free state.

During creation of the mScarlet family, the fluorescence lifetime was included in the screening process, because the authors aimed to optimize the QY linked to the lifetime ^16,17,22^. The resulting mScarlet FPs, with lifetimes of 3.9 ns for mScarlet and 4.0 for mScarlet3 (records for red FPs), could provide a good basis for a biosensor with a lifetime contrast. Other mScarlet variants could be considered as well, due to their maturation speed and lifetime, namely mScarlet-I, mScarlet-I3 and mScarlet-A220 (an intermediate variant of mScarlet3). We have previously developed two calcium sensors with a lifetime contrast of > 1 ns, Tq-Ca-FLITS^21^ and G-Ca-FLITS^23^. Both are based on mTurquoise2, an FP with a fluorescence lifetime of 4.0 ns ^24^. Therefore, we aimed to use mScarlet in a similar fashion, to gain a superior bright red calcium sensor showing a lifetime contrast.

## Results

### mScarlet-A220 and mScarlet-I based sensors

Based on the successful design of Tq-Ca-FLITS, we created mScarlet-A220 (ScA220) and mScarlet-I (ScI) candidate sensors. A circular permutated (cp) mScarlet is flanked by M13 and CaM, with linker sequences in between, identical to the linker sequences in R-GECO1. The mScarlet barrel was opened at positions 144–148 in the 7^th^-strand to find a sensitive spot. This led to 10 variants (**Figure S1**). A dummy sensor was created as a control: a normal mScarlet-A220 flanked by M13 and CaM. The sensors were expressed in bacteria and crudely isolated by a bacterial lysis protocol developed previously ^23^. The protocol allows expression and isolation of 96 variants in parallel and uses a mild lysis buffer with 2% deoxycholic acid (DOC). The performance of the sensors was evaluated in this bacterial lysate by measuring the fluorescence lifetime after addition of calcium and after removal of calcium by the chelator EDTA. Sensor variants starting with residue 147 (cp147) of ScI or ScA220 were the brightest, although still very dim, and the only variants with sufficient intensity to determine the fluorescence lifetime (**Table 1**). However, virtually no lifetime difference was observed between low and high calcium levels. The dummy sensor seems to show an unexpected, very small lifetime difference of 0.06 ns. Therefore, we are cautious to regard such small lifetime changes as “true” changes.

**Table 1:** Lifetime changes of mScarlet-I, mScarlet-A220, mScarlet-I3 and mScarlet-3 based variants. Fluorescence lifetime was measured in bacterial lysate in the presence (0.1 mM CaCl_2_) and absence (9.5 mM EDTA) of calcium, in *n* isolates at room temperature. Reported are the absolute phase (τ_ϕ_) and modulation lifetime (τ_M_) in the presence of calcium, and the difference in phase and modulation lifetime (Δτ_ϕ_ and Δτ_M_) between the two conditions (+Ca^2+^ minus −Ca^2+^), including standard deviation [sd] if applicable. Only sensor variants that showed sufficient fluorescence to determine a fluorescence lifetime are listed in this table. The total number of created variants is indicated in the category headers.

| Internal number | Systematic name | $\tau_\phi$ +Ca<sup>2+</sup><br>[sd] (ns) | $\tau_M$ +Ca<sup>2+</sup><br>[sd] (ns) | $\Delta\tau_\phi$<br>[sd] (ns) | $\Delta\tau_M$<br>[sd] (ns) | n |
| --- | --- | --- | --- | --- | --- | --- |
| *First mScarlet-I and mScarlet-A220 variants (10 in total)* |  |  |  |  |  |  |
| 1632 | cp147mScarlet-I-GECO1 | 1.98 [0.00] | 2.35 [0.01] | 0.03 [0.00] | 0.00 [0.01] | 2 |
| 5304 | cp147mScarlet-A220-GECO1 | 2.47 [0.01] | 2.80 [0.01] | 0.03 [0.01] | 0.05 [0.06] | 2 |
| *Additional mScarlet-I and mScarlet-A220 variants: deletions, mutations, other positions of cp (21 in total)* |  |  |  |  |  |  |
| 5309 | cp147del146-145-CL-mScarlet-A220-GECO | 2.77 [0.07] | 2.96 [0.06] | -0.21 [0.01] | -0.17 [0.00] | 10 |
| 5310 | cp162del161-CL-mScarlet-A220-GECO | 2.43 [0.03] | 2.77 [0.02] | -0.09 [0.03] | -0.14 [0.01] | 2 |
| 5340 | cp148del147mScarlet-A220 GECO | 1.68 [0.00] | 2.12 [0.01] | 0.10 [0.01] | 0.04 [0.05] | 2 |
| 5344 | cp148del147-KL-mScarlet-A220-GECO | 2.24 [0.00] | 2.76 [0.00] | 0.05 [0.00] | 0.05 [0.01] | 2 |
| *mScarlet3 and mScarlet-I3 variants, based on variant 5309 (12 in total)* |  |  |  |  |  |  |
| 5496 | cp147del145-146_Scl3-CL-GECO | 2.51 [0.06] | 2.72 [0.05] | -0.10 [0.01] | -0.03 [0.01] | 14 |
| 5499 | cp148del146-147_Sc3-CL-GECO | 1.38 | 1.73 | -0.21 | -0.19 | 1 |
| 5505 | cp147del145-146_Sc3-CL-GECO | 3.06 [0.01] | 3.15 [0.01] | -0.04 [0.00] | -0.02 [0.00] | 3 |
| *mScarlet3 and mScarlet-I3 variants, screening of mixed PCR (100 new possibilities in total)* |  |  |  |  |  |  |
| 5511 | cp144ins144-RGL/CL-Scl3-GECO | 3.20 [0.03] | 3.30 [0.05] | 0.35 [0.02] | 0.30 [0.01] | 12 |
| 5512 | cp146del145-RGL/CL-Scl3-GECO | 2.54 | 2.77 | -0.21 | -0.13 | 1 |
| *mScarlet3 and mScarlet-I3 variants, additional design (3 in total)* |  |  |  |  |  |  |
| 5515 | cp144ins144-RGL/CL-Sc3-GECO | 3.57 | 3.68 | 0.25 | 0.19 | 1 |
| *Control: dummy sensor* |  |  |  |  |  |  |
| 5313 | cp1mScA220-GECO | 3.90 [0.06] | 3.87 [0.05] | 0.06 [0.01] | 0.06 [0.01] | 10 |

Inspired by the influence of I162, L166 and R198 on the fluorescence lifetime of mScarlet, we created cp162, cp166 and cp198 variants. In addition, copies of the red fluorescent calcium sensors CH-GECO2.1, K-GECO1 and R-GECO1 were made, but with the FP replaced by ScA220 or ScI. Essential mutations and deletions were carefully included in the designs, namely: mutations M164K and T146S in the R-GECO copy, and changes of the linker sequences to match CH-GECO2.1 or K-GECO1. We also included cp162, cp166 and cp198 variants with the linkers of CH-GECO2.1. In total 21 variants were created, including some variants with point mutations (**Figure S1**). The performance of all candidates was evaluated in bacterial lysate as before, in presence and absence of calcium (**Table 1**). Four of the variants were sufficiently bright to determine the lifetime from, and three of those showed a lifetime contrast, namely the CH-GECO2.1 mimic (Δτ_ϕ_ = −0.21 ns and Δτ_M_ = −0.17 ns, number 5309), a cp148 variant very similar to R-GECO1 (Δτ_ϕ_ = 0.10 ns and Δτ_M_ = 0.04 ns, number 5340) and the cp162 variant with the CH-GECO2.1 linker sequence (Δτ_ϕ_ = −0.09 ns and Δτ_M_ = −0.14 ns, number 5310). We choose to further develop the CH-GECO2.1 mimic (referred to as 5309 from now on), as it was visually the brightest of the three and had the largest lifetime contrast.

### Initial mScarlet3 and mScarlet-I3 based sensors

The evolution of mScarlet to the even faster maturating mScarlet3, motivated us to replace mScarlet-A220 in 5309 with mScarlet3 or mScarlet-I3. Also, several other variations with mScarlet3 or mScarlet-I3 were created, with deletions or insertions with respect to 5309 and/or with different linker sequences. This led to 12 variants in total (**Figure S2**). Three of these variants were bright enough to determine the fluorescence lifetime from, and two showed a lifetime difference in bacterial lysate between presence and absence of calcium (**Table 1**, **Figure 1**). Unfortunately, the variant with the largest change (Δτ_ϕ_ = −0.21 ns and Δτ_M_ = −0.19 ns, number 5499) was very dim, while the visually substantially brighter variant showed a smaller lifetime change (Δτ_ϕ_ = −0.10 ns and Δτ_M_ = −0.03 ns, number 5496). This visually brighter variant is identical to 5309 but with mScarlet-I3 as FP.

**Figure 1:**
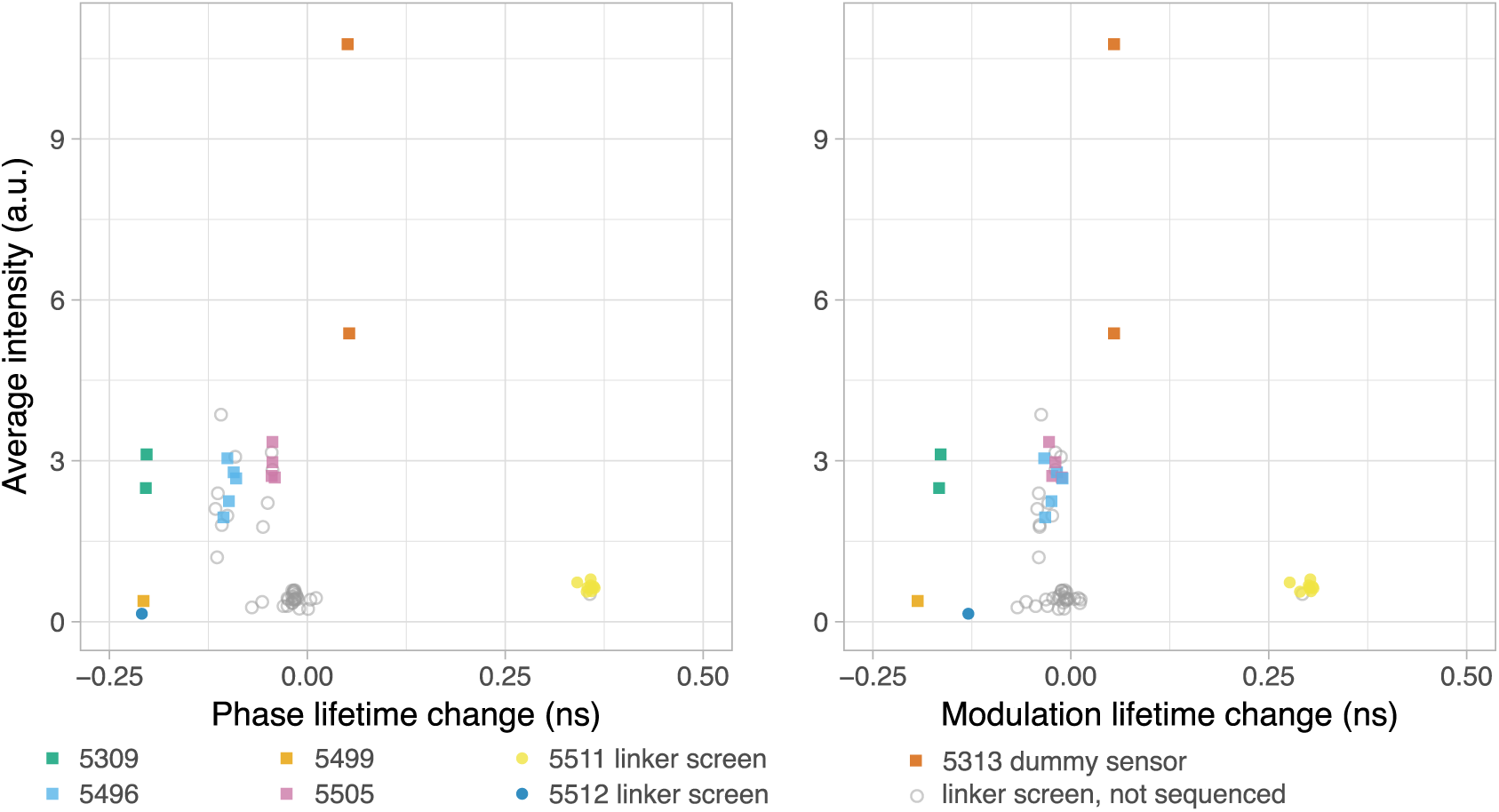
Screening of mSc3 and mSc-I3 sensor variants from a library containing all possible variants with the designed primers. The fluorescence lifetime was measured in bacterial lysate in the presence (0.1 mM CaCl_2_) and absence (9.5 mM EDTA) of calcium at room temperature. The difference in phase and modulation lifetime between the two conditions (+Ca^2+^ minus −Ca^2+^) is plotted against the average intensity of the lysates in the two conditions. Each point represents the measurement of a single lysate originating from a single colony. Variants indicated by a cube were rationally designed previously, variants indicated by a dot are newly found in the screen and variants indicated by a gray circle were not sequenced.

So far, all sensor variants were created in a similar manner: a cpFP was created by PCR using a construct with two FPs in tandem as template. In the primers were sequences included that would become the linkers between the FP and the M13 or CaM. Finally, the PCR product and a backbone containing CaM and M13 were digested and ligated. As the candidate sensors so far were either very weak or lacked a lifetime contrast, we decided to use all primers at once in a single PCR (one for mScarlet3 and one for mScarlet-I3) to create new combinations. In total 112 variants were expected, of which a 100 new variants.

We screened 1180 colonies and picked 60 red fluorescent variants from bacterial plates. These variants were screened in bacterial lysate, looking for variants with a large lifetime contrast and high average brightness. The bacterial lysates of 53 variants showed sufficient red fluorescence to determine the lifetime from, in presence and absence of calcium. A plot of the average intensity of the two states versus the lifetime change shows several clusters. Of each interesting cluster, the sequence of several individuals was determined (**Figure 1, Figure S2**). The sequencing results align very well with the observed clusters, so we were confident that all individual measurements in a cluster belong to the same variant.

Two interesting variants were found with a low intensity, but with lifetime changes of Δτ_ϕ_ = 0.35 ns and Δτ_M_ = 0.30 ns (number 5511), and Δτ_ϕ_ = −0.21 ns and Δτ_M_ = −0.13 ns (number 5512). Some variants were retrieved in the screen that we designed earlier. Based on the results of this screen, three more variants were designed, of which one was fluorescent enough to determine the lifetime change from. This was substantial (Δτ_ϕ_ = 0.25 ns and Δτ_M_ = 0.19 ns, number 5515, **Figure 2**), but less than variant 5511 retrieved from the screen. Notably, of all sensor variants, the CH-GECO copy with mScarlet-I3 as FP (number 5496) was the brightest visually on plate and in bacterial lysate.

**Figure 2:**
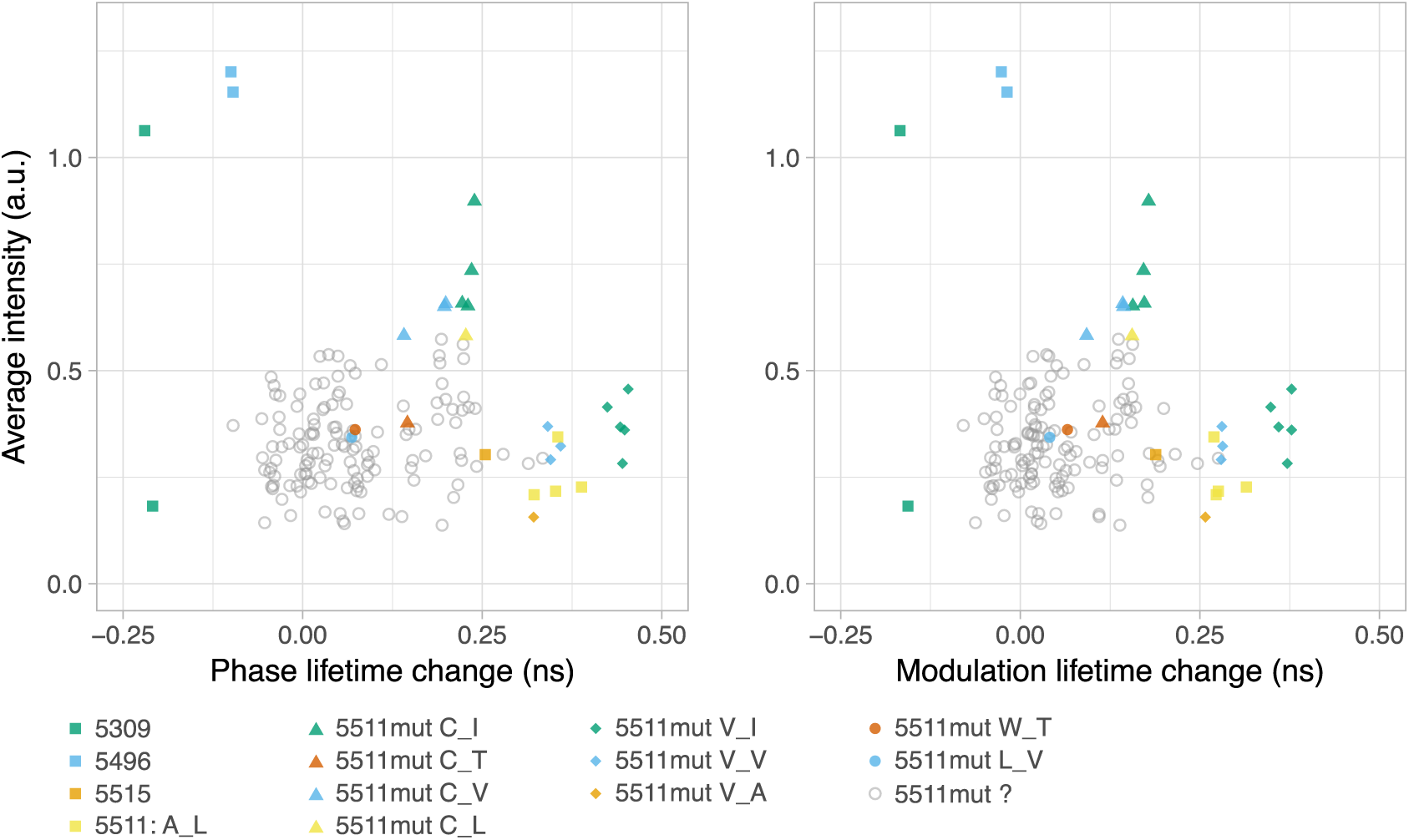
Mutagenesis of sensor variant 5511 on positions A30 and L266. The fluorescence lifetime was measured in bacterial lysates in the presence (0.1 mM CaCl_2_) and absence (9.5 mM EDTA) of calcium at room temperature. The difference in phase and modulation lifetime between the two conditions (+Ca^2+^ minus −Ca^2+^) is plotted against the average intensity of the lysates in the two conditions. Each point represents the measurement of a single lysate originating from a single colony. Colors and shapes are indicative of the mutations, except for variants indicated by a cube: these were rationally designed previously or found in the previous screen. Mutations are indicated by letters in the legend, f.e. 5511mut C_I has mutations A30C and L266I. Variants indicated by a gray circle are non-sequenced mutants.

The dummy sensor was included in the testing, both as control and to get a general idea of the brightness of the candidate sensors. However, the intensity between two replicates of the dummy sensor is highly variable, due to differences in expression level in bacteria. Also, the intensity between different replicates of several sensor variants is variable. Therefore, the intensity of the lysate is not a reliable way to screen for brightness, although it allows for a rough comparison. As we did not create a sensor that was both bright and had a relatively large lifetime change, we decided to continue with the brightest Sc-I3 variant (5496) and with the variant with the largest lifetime change (5511).

### Mutagenesis of red sensor variants

#### Mutagenesis of 5511

Variant 5511 was screened for beneficial mutations at position A30, right after the first linker sequence, and at position L266 in the second linker sequence (sensor numbering). Over 5000 colonies were screened on agar plates for red fluorescence, of which 167 were selected for screening in bacterial lysate (**Figure 2, Table S1**). Of these variants, 153 were bright enough to determine the fluorescence lifetime from and 22 interesting variants were selected for sequencing.

The variant with the largest lifetime contrast was 5511mut V_I (Δτ_ϕ_ = 0.44 ns and Δτ_M_ = 0.37 ns, V_I standing for mutations A30V and L266I), followed by 5511mut V_V (Δτ_ϕ_ = 0.35 ns and Δτ_M_ = 0.28 ns) and the unmutated 5511 (Δτ_ϕ_ = 0.35 ns and Δτ_M_ = 0.30 ns). When looking at the average brightness of the lysates in the calcium-free and -bound situation, variants with mutation A30C perform better (5511mut C_I, 5511mut C_V and 5511mut C_L), but at the expense of the lifetime contrast. All mutants found in this screen, showed a lower intensity than the 5496 variant.

#### Mutagenesis of 5496

Variant 5496 was also subjected to mutagenesis of two amino acids, namely S30, directly after the first linker, and L265 in the second linker (sensor numbering). After mutagenesis, a total of ~ 10, 000 colonies were screened on agar plates, of which 176 red fluorescent ones were selected for screening in bacterial lysate (**Figure 3, Table S1**). Here, the lifetime difference between calcium-free and calcium-bound could be determined of all except one variant. We selected 23 variants of different interesting phenotypes (absolute lifetime and/or lifetime change) for sequencing. Hence, not all improved variants were sequenced. The screen was repeated he next day, with 176 new colonies picked from the same agar plates. Of all variants, the lifetime change could be determined. 20 interesting variants were sequenced, and similar mutants were found as on the first day.

**Figure 3:**
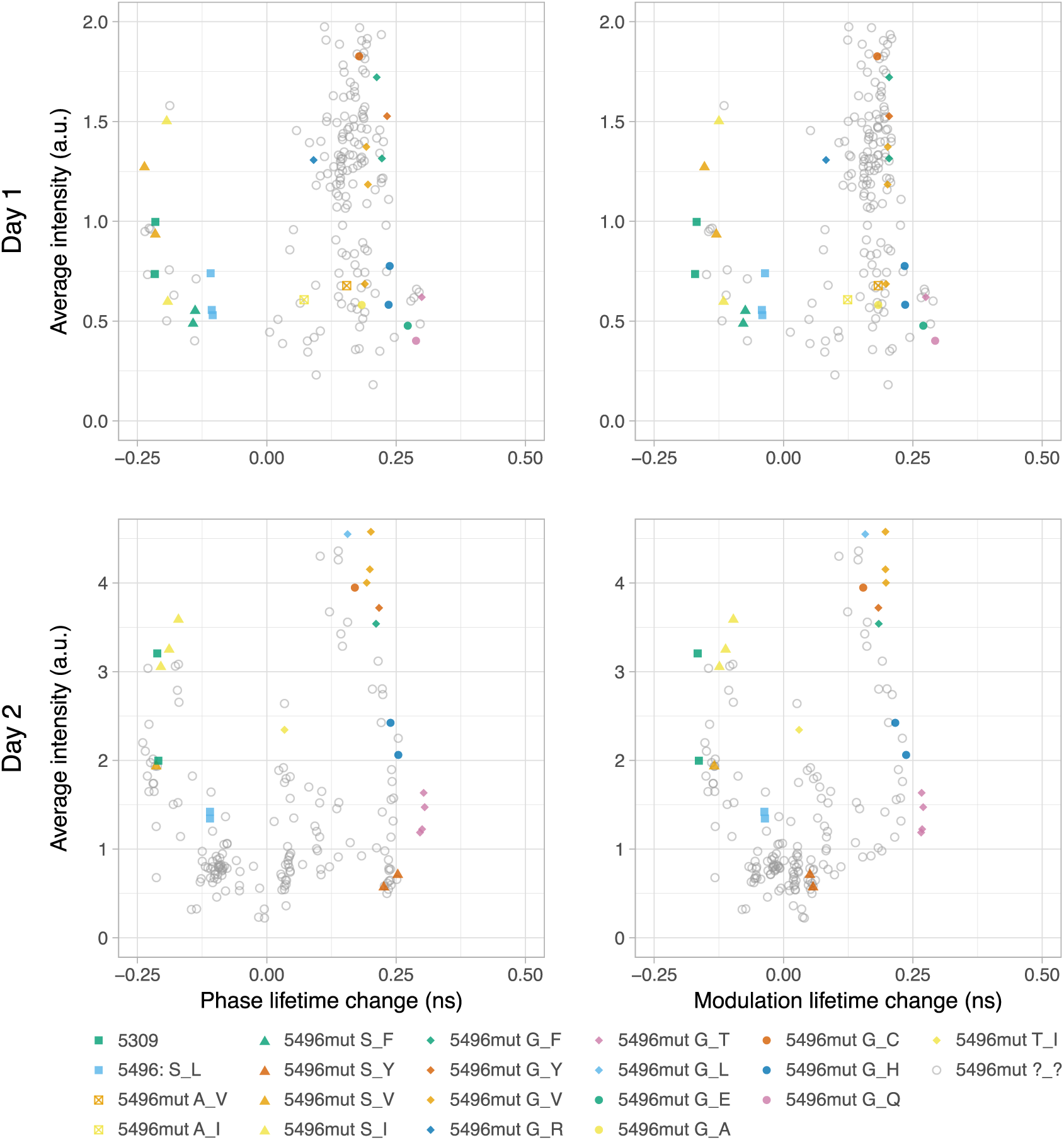
Mutagenesis of sensor variant 5496 on positions S30 and L265. The fluorescence lifetime was measured in bacterial lysates in the presence (0.1 mM CaCl_2_) and absence (9.5 mM EDTA) of calcium at room temperature. The difference in phase and modulation lifetime between the two conditions (+Ca^2+^ minus −Ca^2+^) is plotted against the average intensity of the lysates in the two conditions. The screen was performed on two days and data of these days are plotted separately. Each point represents the measurement of a single lysate originating from a single colony. Colors and shapes are indicative of mutations, except for variants indicated by a cube: these were rationally designed previously. Mutations are indicated by letters in the legend, f.e. 5496mut G_Y has mutations S30G and L265Y. Mutant sensors indicated by a gray circle were not sequenced.

All 11 variants with a L265G mutation show a positive lifetime change (calcium present minus calcium absent), while three out of four variants with a S at this position (as in the parent 5496) show a negative lifetime change. The variants with the largest lifetime changes are 5496mut G_Q (Δτ_ϕ_ = 0.29 ns and Δτ_M_ = 0.29 ns, mutations S30G and L265Q), 5496mut G_T (Δτ_ϕ_ = 0.30 ns and Δτ_M_ = 0.27 ns) and 5496mut G_E (Δτ_ϕ_ = 0.27 ns and Δτ_M_ = 0.27 ns). When looking at the average brightness of the lysates in the screen, the variants with the highest lifetime change are less bright compared to other variants. The brightest variants were 5496mut G_L, 5496mut G_F, 5496mut G_Y, 5496mut G_V and 5496mut G_C. No variant in this screen showed a comparable lifetime contrast to variant 5511mut V_I from the previous mutagenesis.

#### Mutagenesis of position R198

We hypothesized that the chromophore in mScarlet-I3 fluorescent protein is too rigid to be sensitive to conformational changes, and that disrupting the rigidity might have a beneficial effect on the dynamic range. It is known from the engineering of mScarlet that R198 has a strong stabilizing effect on the chromophore ^16^. Therefore, a directed R198I mutagenesis was performed in the variants 5496mut G_Y, 5496mut S_V, 5511mut V_I and 5511mut C_I. Unfortunately, this reduced the fluorescence in bacterial lysates to a level that was too low for determination of the fluorescence lifetime, and we did not continue with these variants.

### A ratiometric plasmid for comparison of intensities

So far, no sensor variant with both a high brightness and a large lifetime contrast was developed. Therefore, we wanted to improved our screening approach by including a robust way of screening for the fluorescence brightness of sensor variants, that is not influenced by bacterial growth or protein expression. To this end, a dual expression ratio plasmid was designed, named pFR (for Franka Ratio). It contains mTurquoise2 (mTq2) followed by an anti-FRET linker, a P2A sequence and a red calcium sensor variant (**Figure S3**). The plasmid is suitable for expression both in bacteria (rhamnose promotor) and mammalian cells (CMV promotor). In bacteria, the anti-FRET linker reduces energy transfer from the mTq2 to the red sensor. In mammalian cells, the P2A sequence is cleaved, resulting in two separate proteins. On the N-terminal side, the fixed mTq2 has a His-tag for protein isolation. Within a mScarlet calcium sensor, the cpFP can be replaced using MluI and SacI restriction sites. The calcium sensor can be completely replaced using the BamHI and HindIII restriction sites. The mTq2, anti-FRET linker and P2A sequence can be removed by digestion with NheI, creating a plasmid with only a His-tagged sensor, named pFPO, for plasmid Franka Protein Only.

It was verified if the pFR plasmid is suitable for screening of libraries of sensors in bacteria, and comparison of the intensity of sensors in HeLa cells. First, we compared the lifetime of cyan fluorescence on agar plates between bacteria expressing mTq2 or mTq2 attached via the anti-FRET linker to the dummy sensor. A clear difference in lifetime was measured: addition of the dummy sensor reduces the modulation lifetime of mTq2 from 4.0 ns to below 3.0 ns, indicative of Förster Resonance Energy Transfer (FRET) (**Figure S4A**). We observed a variation in the lifetime for both constructs, which was related to the position of the colony to the center of the microscopy images. A second pFR plasmid was also tested, with mScI as fixed plasmid instead of mTq2, and with a green calcium sensor (Tq-Ca-FLITS_T203Y, an intermediate of G-Ca-FLITS^23^, instead of a red one. The same results were observed in bacterial colonies: the fluorescence lifetime of Tq-Ca-FLITS_T203Y was reduced more than 1 ns when mScI was present (**Figure S4B**), with a positional dependent lifetime. The reduced lifetimes of mTq2 and Tq-Ca-FLITS_T203Y in the presence of mSc, are likely caused by bystander FRET due to the high concentration of proteins in the bacterial cytoplasm.

Second, we made a crude protein isolate from bacteria expressing mTq2, with and without the red dummy sensor attached. The fluorescence lifetime of the mTq2 was compared between the lysates, and between addition of CaCl_2_ or EDTA. There was no difference in the lifetime between the isolates, or between the different concentrations of calcium (**Table 2**). Next, the dummy sensor was photobleached to almost 60% of its original intensity. This did not affect the fluorescence lifetime of mTq2. The same procedure was applied to the second ratio plasmid, comparing Tq-Ca-FLITS_T203Y to mScI-anti-FRET-Tq-Ca-FLITS_T203Y. In all situations the lifetime of Tq-Ca-FLITS_T203Y was reduced by mScI; however, the lifetime contrast of Tq-Ca-FLITS_T203Y between the calcium-free and calcium-saturated state remains almost the same (Δτ_ϕ_ = −1.1 vs −1.2 ns and Δτ_M_ = −1.2 vs −1.3 ns). Photobleaching of mScI to about 60% had no significant effect on the lifetime.

**Table 2:** Influence of an RFP on the fluorescence lifetime of mTq2 and Tq-Ca-FLITS_T203Y in bacterial lysate. The fluorescence phase (τ_ϕ_) and modulation lifetime (τ_M_) were measured in bacterial lysate at room temperature, in presence (0.1 mM CaCl_2_) or absence (9.5 mM EDTA) of calcium, and before or after photobleaching of the RFP to 60% of its original intensity. The standard deviation (sd) over the full field of view of the microscope indicated. *The same bleaching pulse was applied as for the samples with RFP.

| Protein | bleached RFP | +Ca<sup>2+</sup> |  |  |  | -Ca<sup>2+</sup> |  |  |  |
| --- | --- | --- | --- | --- | --- | --- | --- | --- | --- |
| | | $\tau_\phi$ (ns) | sd | $\tau_M$ (ns) | sd | $\tau_\phi$ (ns) | Sd | $\tau_M$ (ns) | sd |
| mTq2 | no | 4.29 | 0.04 | 4.27 | 0.04 | 4.27 | 0.04 | 4.26 | 0.04 |
| mTq2 | yes* | 4.25 | 0.05 | 4.25 | 0.04 | 4.27 | 0.04 | 4.26 | 0.04 |
| mTq2 - cp1mScA220-GECO | no | 4.26 | 0.05 | 4.24 | 0.04 | 4.20 | 0.05 | 4.19 | 0.04 |
| mTq2 - cp1mScA220-GECO | yes | 4.25 | 0.05 | 4.23 | 0.04 | 4.17 | 0.05 | 4.19 | 0.05 |
| Tq-Ca-FLITS_T203Y | no | 2.21 | 0.02 | 2.85 | 0.03 | 3.41 | 0.04 | 4.10 | 0.04 |
| Tq-Ca-FLITS_T203Y | yes* | 2.18 | 0.03 | 2.81 | 0.04 | 3.34 | 0.04 | 4.05 | 0.04 |
| mScl - Tq-Ca-FLITS_T203Y | no | 2.05 | 0.02 | 2.45 | 0.03 | 3.18 | 0.03 | 3.74 | 0.03 |
| mScl - Tq-Ca-FLITS_T203Y | yes | 2.05 | 0.02 | 2.45 | 0.04 | 3.15 | 0.03 | 3.73 | 0.04 |

Third, we expressed both ratio plasmids and the controls (without mScI or the dummy sensor) in HeLa cells and investigated fluorescence lifetime of mTq2 and Tq-Ca-FLITS_T203Y (**Figure 4A, Table S2**). Again, for both ratio plasmids the presence of the RFP led to a small reduction in the lifetime. Photobleaching of the RFP to 22% of its original intensity did not have a significant effect. In addition, we added ionomycin (5 µg/mL) and extra calcium (5 mM) to raise the calcium level in the cells, in order to saturate the (dummy) calcium sensor (**Figure 4B, Table S2**). Only a lifetime response is seen for the constructs with Tq-Ca-FLITS_T203Y. Here, the lifetime contrast is a little reduced in the ratio plasmid with mScI compared to normal Tq-Ca-FLITS_T203Y (Δτ_ϕ_ = −0.60 ns and Δτ_M_ = −0.72 ns versus Δτ_ϕ_ = −0.67 ns and Δτ_M_ = −0.78 ns), but this effect was not significant (**Table S2**).

**Figure 4:**
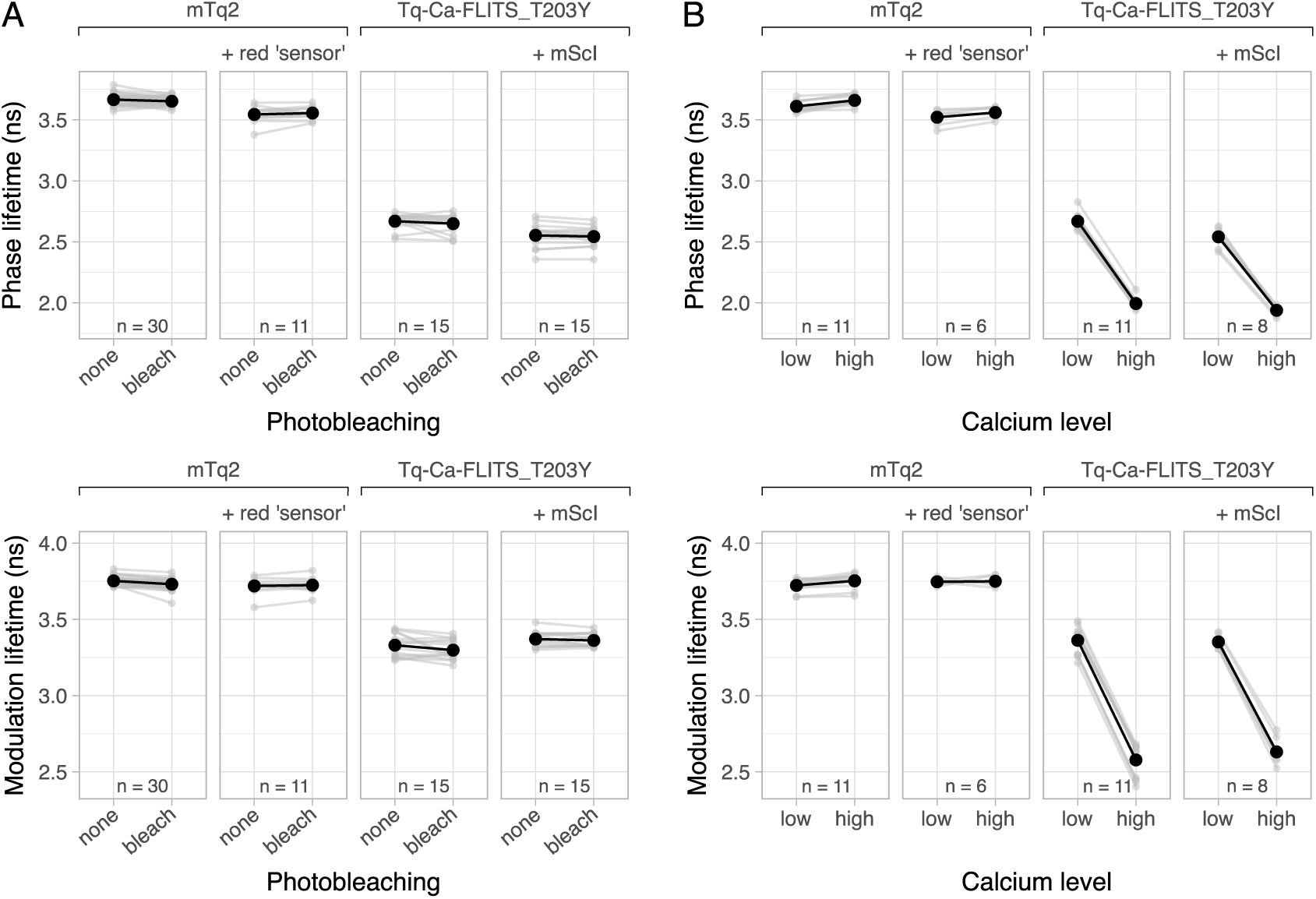
Influence of the calcium concentration or photobleaching of the RFP component on the phase and modulation lifetime of mTq2 and Tq-Ca-FLITS_T203Y in HeLa cells. **A)** The fluorescence lifetimes were measured of the mTq2 or the Tq-Ca-FLITS_T203Y component of the indicated constructs, at 37 °C, before and after photobleaching of the RFP to 22% of its original intensity. The same bleaching pulse was applied to samples without an RFP. For each individual cell the phase (Δτ_ϕ_) and modulation lifetime changes (Δτ_M_) are calculated. Individual cells are indicated in gray, with lines connecting the measurements that belong to a single cell, and with the averages of all cells (n indicated in the plots) indicated in black. **B)** Same as **A**, but measurements were taken before and after addition of 5 µg/mL ionomycin and 5 mM CaCl_2_ (low and high calcium level), instead of before and after photobleaching.

In summary, both ratio plasmids show evidence of bystander FRET in bacterial colonies on plate, indicated by the discrepancy in the lifetimes of mTq2 and Tq-Ca-FLITS_T203Y in the presence and absence of mSc. The high concentration of proteins in the bacterial cytosol allows for FRET between the green/cyan FP in one chain and the RFP in another chain, reducing the lifetime. The bacterial lysate is a diluted version of the bacterial cytoplasm, where chains are too far apart for bystander FRET to occur. Indeed, the the fluorescence lifetime of mTq2 and Tq-Ca-FLITS_T203Y is not influenced by mSc in the bacterial lysate. Due to bystander FRET, screening of red or green calcium sensors based on intensity is not possible in bacterial colonies. However, the pFR plasmid with fixed mTq2 can be used for picking of red sensor variants from agar plates from a mutagenesis library. Here, colonies are chosen based on red fluorescence only, as mTq2 it is not excited and thus cannot contribute to RFP intensity by means of bystander FRET. The pFR plasmid with a fixed mScI can also be used for choosing colonies containing green calcium sensors after a round of mutagenesis, as we expect that bystander FRET levels will be comparable for sensors with only a few mutations, and thus the green fluorescence of these colonies will be equally reduced by FRET. No FRET was observed in either bacterial lysates or HeLa cells, although a small difference in lifetime of mTq2 and Tq-Ca-FLITS_T203Y was measured, but not in the lifetime contrast of the Tq-Ca-FLITS_T203Y sensor. Therefore, the ratio plasmids can be safely used for comparison of lifetime contrast and intensity by ratiometric measurements, but the absolute lifetimes should be evaluated from cells expressing only the sensors.

### Simultaneously improving intensity and lifetime contrast

#### Mutagenesis of 5511mut V_I

The sensor variant with the largest lifetime change so far (5511mut V_I), was subjected to mutagenesis of two new positions in the linkers: residues E29 and G267 (sensor numbering). The new ratiometric plasmid was used for the screen, and a total of ~ 4000 colonies on agar plates were observed. 129 red fluorescent colonies were selected for screening in bacterial lysate. The fluorescence lifetime change between the calcium-bound and -free state was determined for all except 14 variants (**Figure 5, Table S3**). These variants showed a too weak fluorescence in the calcium-free state. In addition, the red fluorescence intensity was measured, and divided over the fluorescence intensity of mTq2 (**Figure 5**).

**Figure 5:**
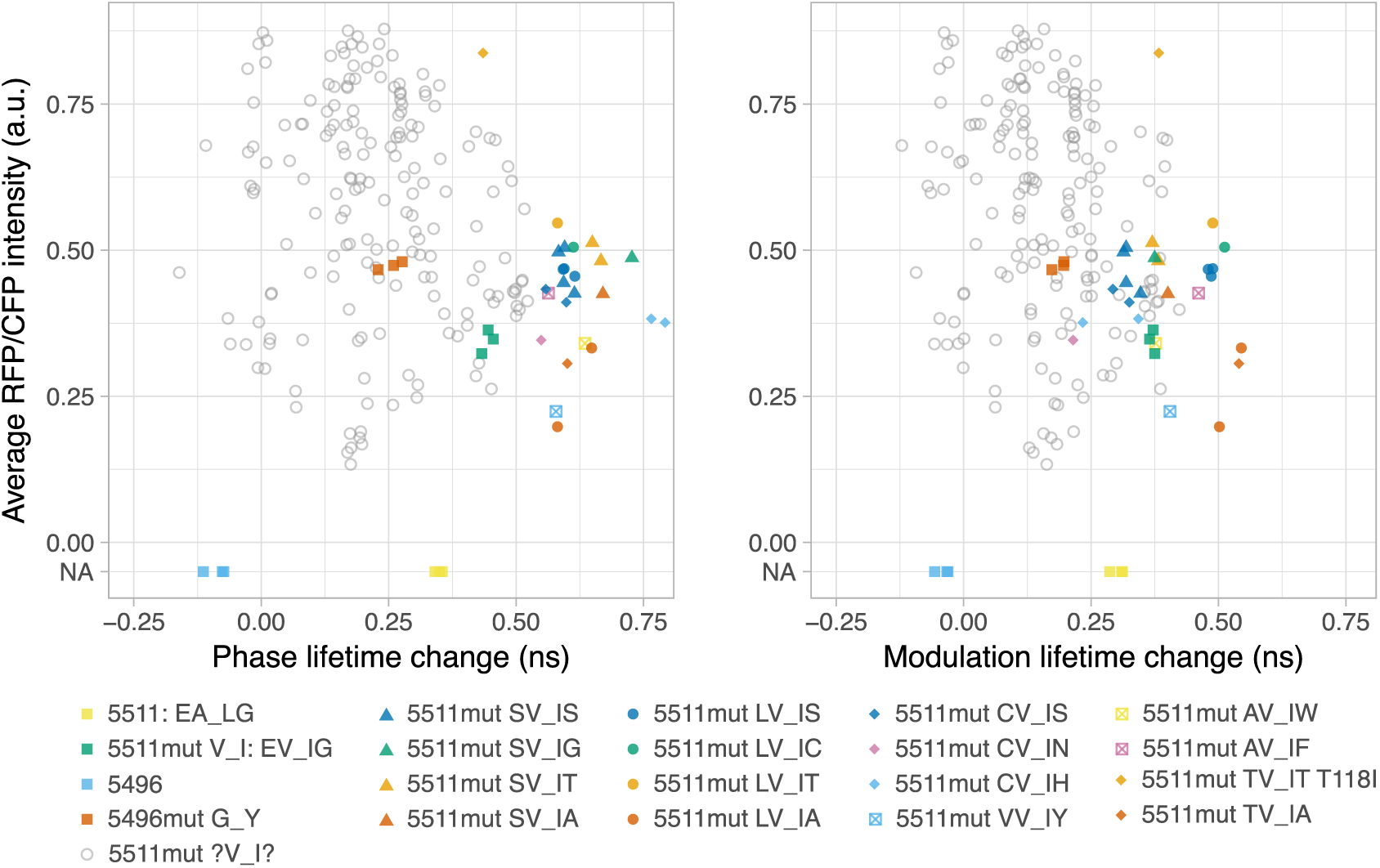
Fluorescence lifetime of sensor variants of 5511mut V_I with mutations on positions E29 and G267. Bacteria expressed simultaneously a red candidate sensor and mTq2 coded on a pFR plasmid. Red and cyan fluorescence, and the fluorescence lifetime of the red candidate sensor were measured in bacterial lysate in the presence (0.1 mM CaCl_2_) and absence (9.5 mM EDTA) of calcium at room temperature. The difference in phase and modulation lifetime (Δτ_ϕ_ and Δτ_M_) between the two conditions (+Ca^2+^ minus −Ca^2+^) are plotted on the x-axis. Red fluorescence was divided over the cyan fluorescence (RFP/CFP) and the average is plotted on the y-axis. Each point represents the measurement of a single lysate originating from a single colony. Colors and shapes are indicative of the mutations as indicated in the legend, f.e. 5511mut LV_IA and has mutations E29L and G267A with respect to 5511mut V_I. Mutant sensors indicated by a gray circle were not sequenced. Of two earlier variants, 5511 and 5496, no RFP/CFP ratio was measured as these were not expressed with pFR.

The majority of the variants showed a smaller lifetime change than the parent 5511mut V_I, and a higher RFP/CFP ratio. The fold-change of the intensity ratio was calculated, and indicated a few mutants with an intensity fold-change of 15 to 20, mainly due to a very low RFP/CFP ratio in the calcium-free state (**Figure 6, Table S3**). Direct comparison of the RFP/CFP intensity of 5496mut G_Y and 5511mut V_I shows that 5496mut G_Y is about 30% brighter compared to 5511mut V_I (**Figure 5**).

**Figure 6:**
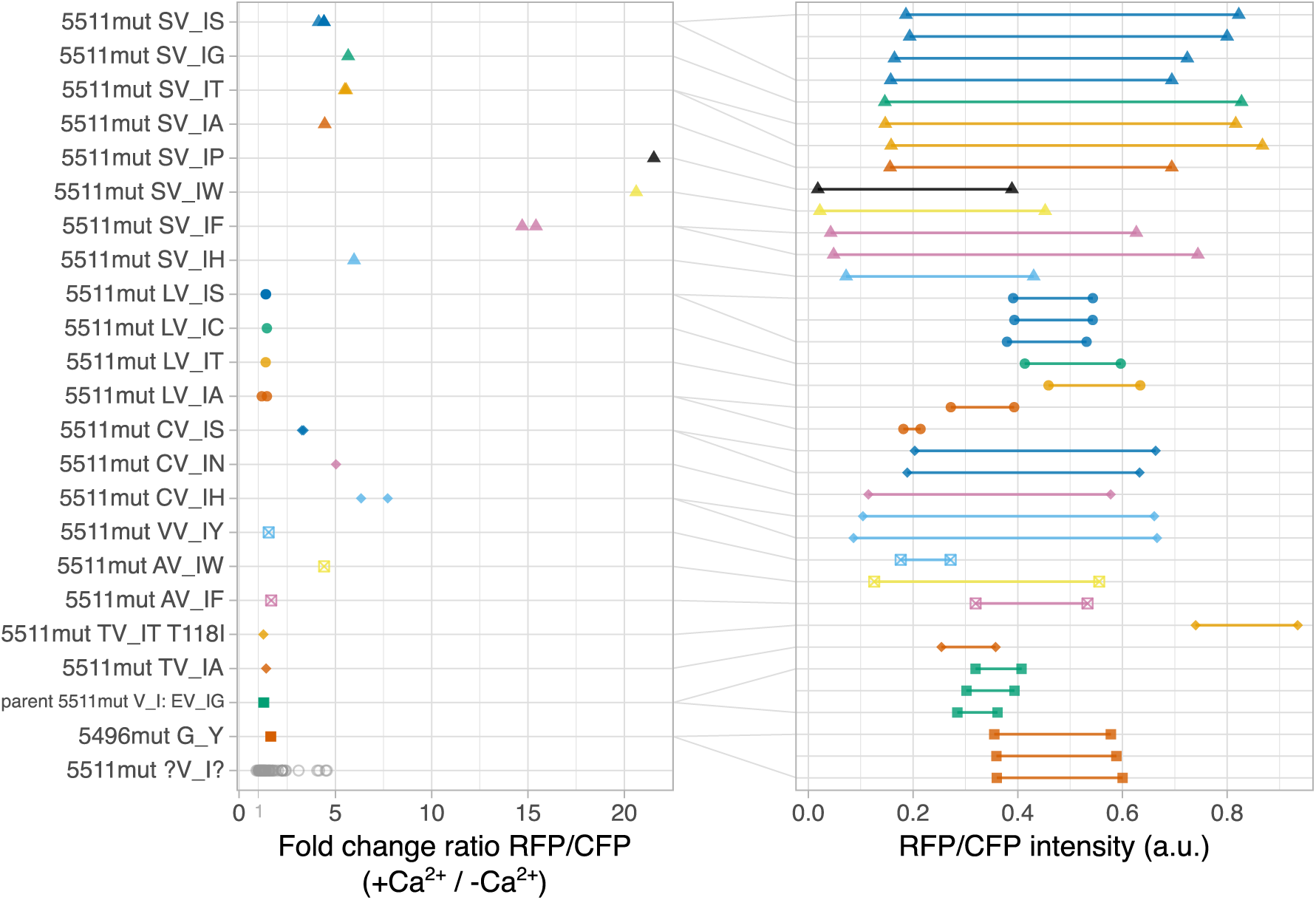
Fluorescence intensities of sensor variants of 5511mut V_I with mutations on positions E29 and G267. Bacteria expressed simultaneously a red candidate sensor and mTq2 coded on a pFR plasmid. Red and cyan fluorescence were measured in bacterial lysate in the presence (0.1 mM CaCl_2_) and absence (9.5 mM EDTA) of calcium at room temperature. Red fluorescence was divided over the cyan fluorescence (RFP/CFP). The fold-changes (+Ca^2+^ / −Ca^2+^) of the ratios are plotted in the left panel, the absolute ratios are plotted in the right panel with a line connecting the two calcium states. Each point in the left panel represents the value of a single lysate originating from a single colony. Colors and shapes are indicative of the mutations as indicated in the legend, f.e. 5511mut LV_IA and has mutations E29L and G267A with respect to 5511mut V_I. Mutant sensors indicated by a gray circle were not sequenced.

We selected 31 variants for sequencing, with either an increased fluorescence lifetime change or a large intensity fold-change. Among the variants with an improved fluorescence lifetime change, mutations to an A, L, S, T or C at position 29 (original E29) were found at least twice. Many different residues were found at position 267, but only A, T and S were present in three variants (original G267). The variants with the largest lifetime contrasts were 5511mut TV_IA (Δτ_ϕ_ = 0.60 ns and Δτ_M_ = 0.54 ns, mutations E29T and G267A), 5511mut LV_IA (Δτ_ϕ_ = 0.61 ns and Δτ_M_ = 0.52 ns), 5511mut LV_IC (Δτ_ϕ_ = 0.61 ns and Δτ_M_ = 0.51 ns) and 5511mut SV_IG (Δτ_ϕ_ = 0.73 ns and Δτ_M_ = 0.37 ns).

The variants with an intensity fold-change between 15 and 20 all had an E29S mutation, combined with either G267P, G267W or G267F. One variant had a similar lifetime change as the parent, but a greatly improved intensity. Sequencing of this variant showed E29T and G267I mutations, alongside an unintended T118I mutation. No E29T plus G267I mutant was found without this extra mutation, so it is unclear whether the T118I mutation is beneficial for the intensity.

#### Mutagenesis of 5496mut G_Y

Also 5496mut G_Y was subjected to mutagenesis of two additional amino acids, namely residue L29 in the first linker and G267 in the second linker, aiming for a brighter variant with increased lifetime contrast. We screened a total of ~ 3500 colonies on agar plates, of which 130 red fluorescent ones were selected for screening in bacterial lysate. The fluorescence lifetime difference between the calcium-bound and -free state was determined for all except one variant, of which the fluorescence in the calcium-free state was too low to determine a lifetime (**Figure 7, Table S3**). Most of the screened mutants showed a larger lifetime change than the parent. Again, also the RFP/CFP intensity ratio was determined for both states and the fold-change was calculated. (**Figure 7**, **Figure 8, Table S3**). About half the variants show a brighter intensity than the parent, and almost all variants show a fold-change between 1 and 2.

**Figure 7:**
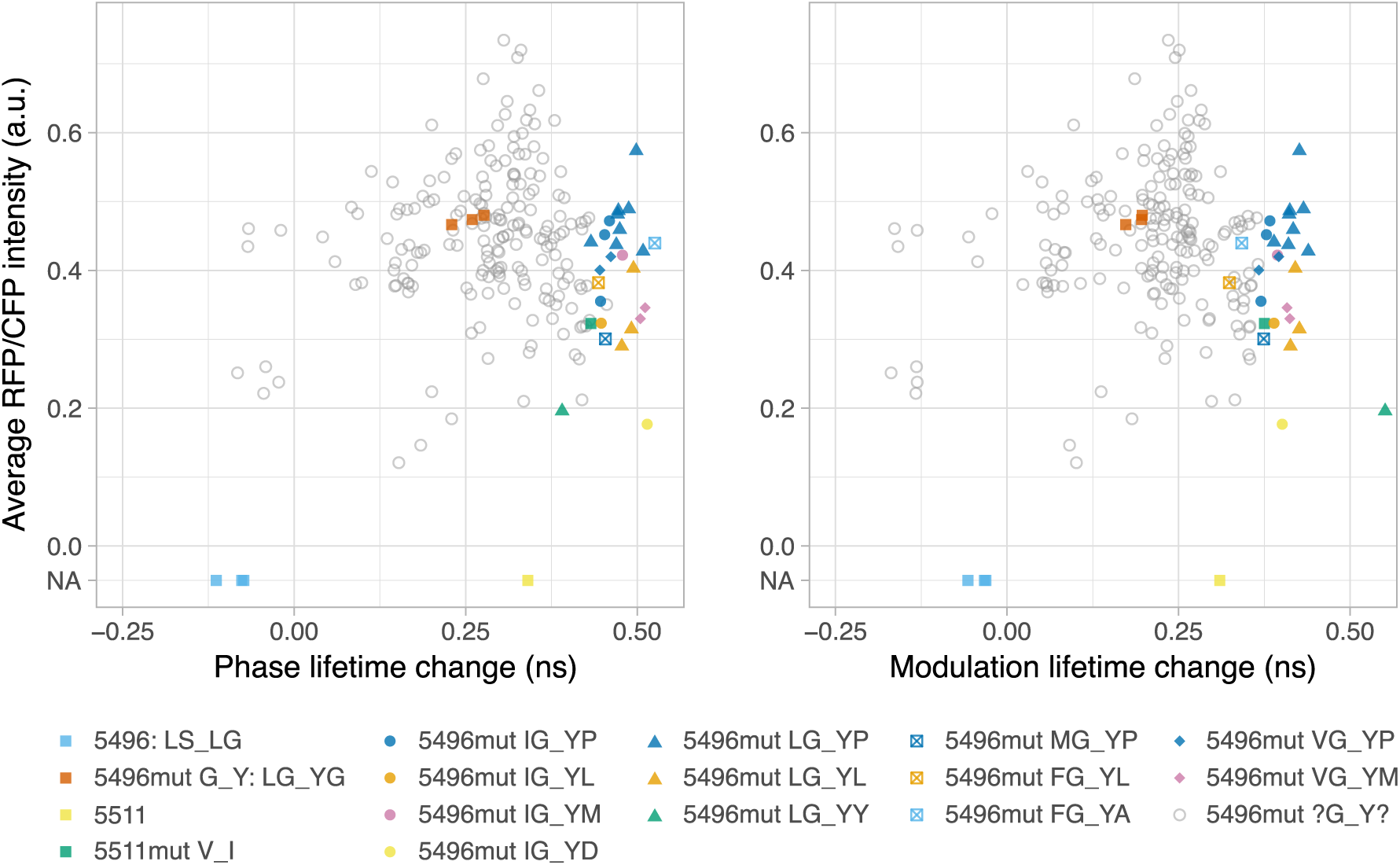
Fluorescence lifetime of sensor variants of 5496mut G_Y with mutations on positions L29 and G266. Bacteria expressed simultaneously a red candidate sensor and mTq2 coded on a pFR plasmid. Red and cyan fluorescence, and the fluorescence lifetime of the red candidate sensors were measured in bacterial lysate in the presence (0.1 mM CaCl_2_) and absence (9.5 mM EDTA) of calcium at room temperature. The difference in phase and modulation lifetime (Δτ_ϕ_ and Δτ_M_) between the two conditions (+Ca^2+^ minus −Ca^2+^) are plotted on the x-axis. Red fluorescence was divided over the cyan fluorescence (RFP/CFP) and the average RFP/CFP ratio of both states is plotted on the y-axis. Each point represents the measurement of a single lysate originating from a single colony. Colors and shapes are indicative of the mutations as indicated in the legend, f.e. 5496mut IG_YP and has mutations L29I and G266P with respect to 5496mut G_Y. Mutant sensors indicated by a gray circle were not sequenced. Of two earlier variants, 5511 and 5496, no RFP/CFP ratio was measured as these were not expressed with pFR

**Figure 8:**
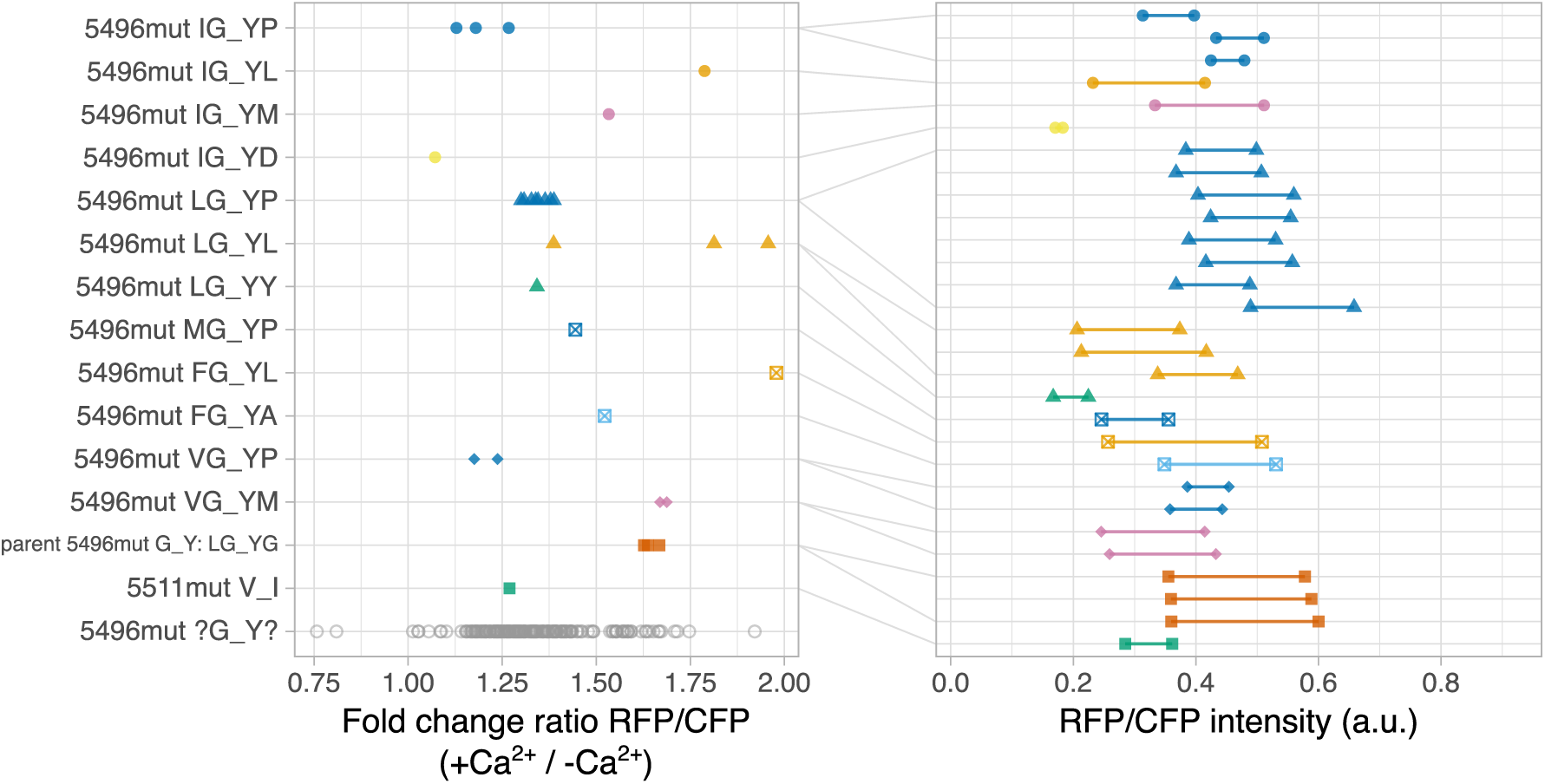
Fluorescence intensities of sensor variants of 5496mut G_Y with mutations on positions L29 and G266. Bacteria expressed simultaneously a red candidate sensor and mTq2 coded on a pFR plasmid. Red and cyan fluorescence were measured in bacterial lysate in the presence (0.1 mM CaCl_2_) and absence (9.5 mM EDTA) of calcium at room temperature. Red fluorescence was divided over the cyan fluorescence (RFP/CFP). The fold-changes (+Ca^2+^ / −Ca^2+^) of the ratios are plotted in the left panel, the absolute ratios are plotted in the right panel with a line connecting the two calcium states. Each point in the left panel represents the value of a single lysate originating from a single colony. Colors and shapes are indicative of the mutations as indicated in the legend, f.e. 5496mut IG_YP and has mutations L29I and G266P with respect to 5496mut G_Y. Mutant sensors indicated by a gray circle were not sequenced.

In total 28 variants that had increased fluorescence lifetime changes compared to the parent were selected for sequencing. Among these, residues V, I, L or F was found at least twice at position L29. At position G266, a mutation to P, L or M was most prevalent among the sequenced variants. The largest lifetime changes were found for 5496mut LG_YY (Δτ_ϕ_ = 0.39 ns and Δτ_M_ = 0.55 ns, mutations L29G and G266Y), 5496mut VG_YM (Δτ_ϕ_ = 0.51 ns and Δτ_M_ = 0.41 ns) and 5496mut IG_YD (Δτ_ϕ_ = 0.51 ns and Δτ_M_ = 0.40 ns).

### Intensity of red calcium sensors in mammalian cells

Overall, the variants generated from 5511mut V_I show the best lifetime contrast, with only a small reduction in intensity compared to 5496mut G_Y mutants. However, all measurements were performed in bacterial lysates. We selected a subset of all mutants in HeLa cells, alongside several published red calcium sensors. The sensors were expressed using the ratiometric plasmid pFR for robust comparison of the intensity. The data was normalized to a control with mCherry (set to 1), and a negative control without red fluorescence (set to 0). The published sensors all show < 18% of the intensity of mCherry in unstimulated cells 24 h after transfection (**Figure 9**). This was improved to < 37% at 48 h post transfection. Ionomycin and extra calcium was added to the cells to increase the intracellular calcium levels to > 5 mM. Upon addition, all sensors show a significantly increased intensity, except CH-GECO2.1.

**Figure 9:**
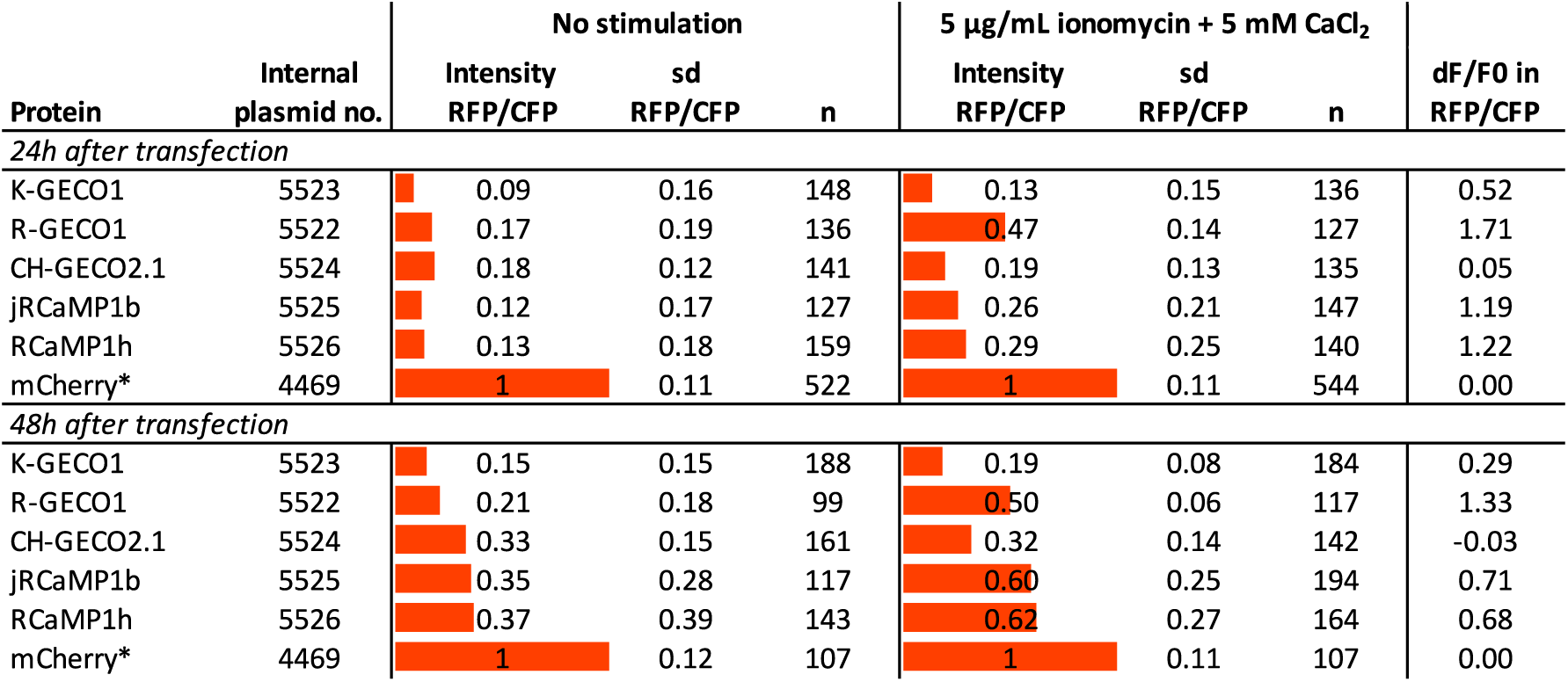
Intensity of published sensors compared to mCherry in HeLa cells. Red and cyan fluorescence was measured in HeLa cells at 37 °C, at both 24- and 48-hours post-transfection with a pFR plasmid coding a red calcium sensor and mTq2. The ratio RFP/CFP was determined for each cell and normalized to mCherry and no RFP present (only mTq2 expression). Averages of all cells (*n)* and the standard deviation (sd) are indicated. The measurements were repeated after addition of 5 µg/mL ionomycin and 5 mM CaCl_2_. The change in RFP/CFP ratio divided over the RFP/CFP ratio before addition (d*F*/*F*_0_) was calculated. *This was measured on another day as all other values in this figure.

Next, mutants from 5511 (mutations at positions A30 and L266) and 5496 (mutations at positions S30 and L265) were evaluated. At 24 h post transfection, all are comparable or weaker in intensity than their parent (**Figure 10**), but the weakest variant 5511mut V_I was still brighter than all published sensors. At 48 h after transfection, 5511 and all its mutants showed an improved intensity, up to twice as much as at 24 h. Barely any significant change in intensity was observed when the intracellular calcium level was raised.

**Figure 10:**
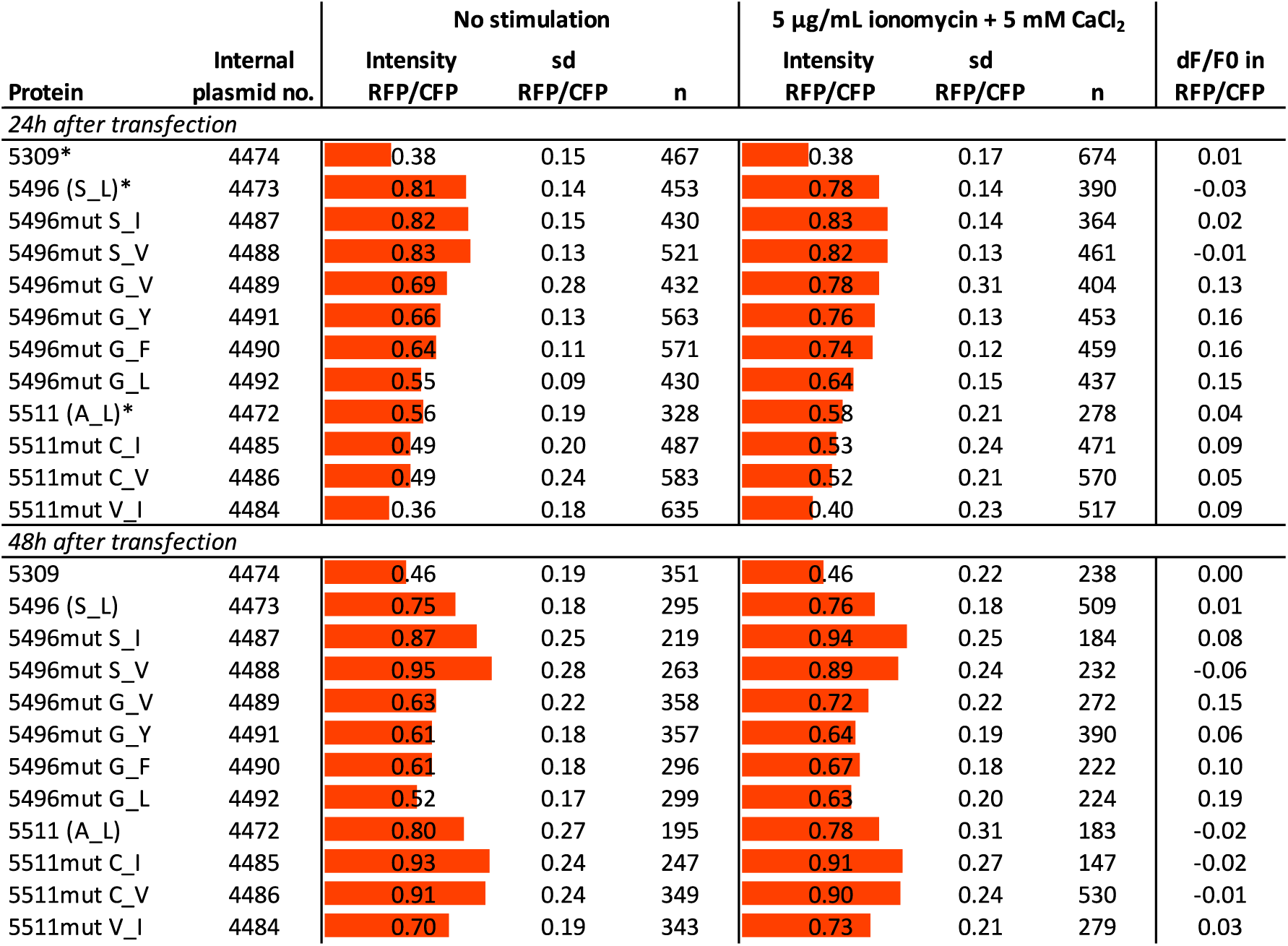
Intensity of 5511 and 5496 mutant sensors in HeLa cells. Red and cyan fluorescence was measured in HeLa cells at 37 °C, at both 24- and 48-hours post-transfection with a pFR plasmid coding a red calcium sensor and mTq2. The ratio RFP/CFP was determined for each cell and normalized to mCherry and no RFP present (only mTq2 expression). mCherry used for normalization was measured on another day. Averages of all cells (*n)* and the standard deviation (sd) are indicated. The measurements were repeated after addition of 5 µg/mL ionomycin and 5 mM CaCl_2_. The change in RFP/CFP ratio divided over the RFP/CFP ratio before addition (d*F/F_0_)* was calculated. Mutants are named after their ancestor and mutations, f.e. 5511mut C_I stems from variant 5511 and has mutations A30C and L266I, and 5496mut G_Y originates from variant 5496 with mutations S30G and L265Y. The original residues at the positions of mutations are indicated at the ancestors 5511 and 5496. *Measured on another day as all other measurements 24 h post transfection in this table.

Finally, the intensity of the mutants of 5511mut V_I (mutations at positions E29 and G267) and 5496mut G_Y (mutations at positions L29 and G266) were compared to mCherry in HeLa cells. All selected variants originating from 5511mut V_I were brighter than their parent, both at 24 h and 48 h post transfection. The selected variants from 5496mut G_Y were either comparable or weaker than their parent. This is as expected based on the intensity in bacterial lysates. When comparing the intensity in HeLa cells with the intensity measured in bacterial lysates, we see a positive correlation within the two mutant groups (**Figure 11, Figure S5**). Raising the intracellular calcium levels with ionomycin at either 24 h or 48 h post-transfection, led to a significant change in intensity in only a few sensors. Notably, the sensors that showed a > 5-fold change in bacterial lysates do not show a significant intensity change in HeLa cells.

**Figure 11:**
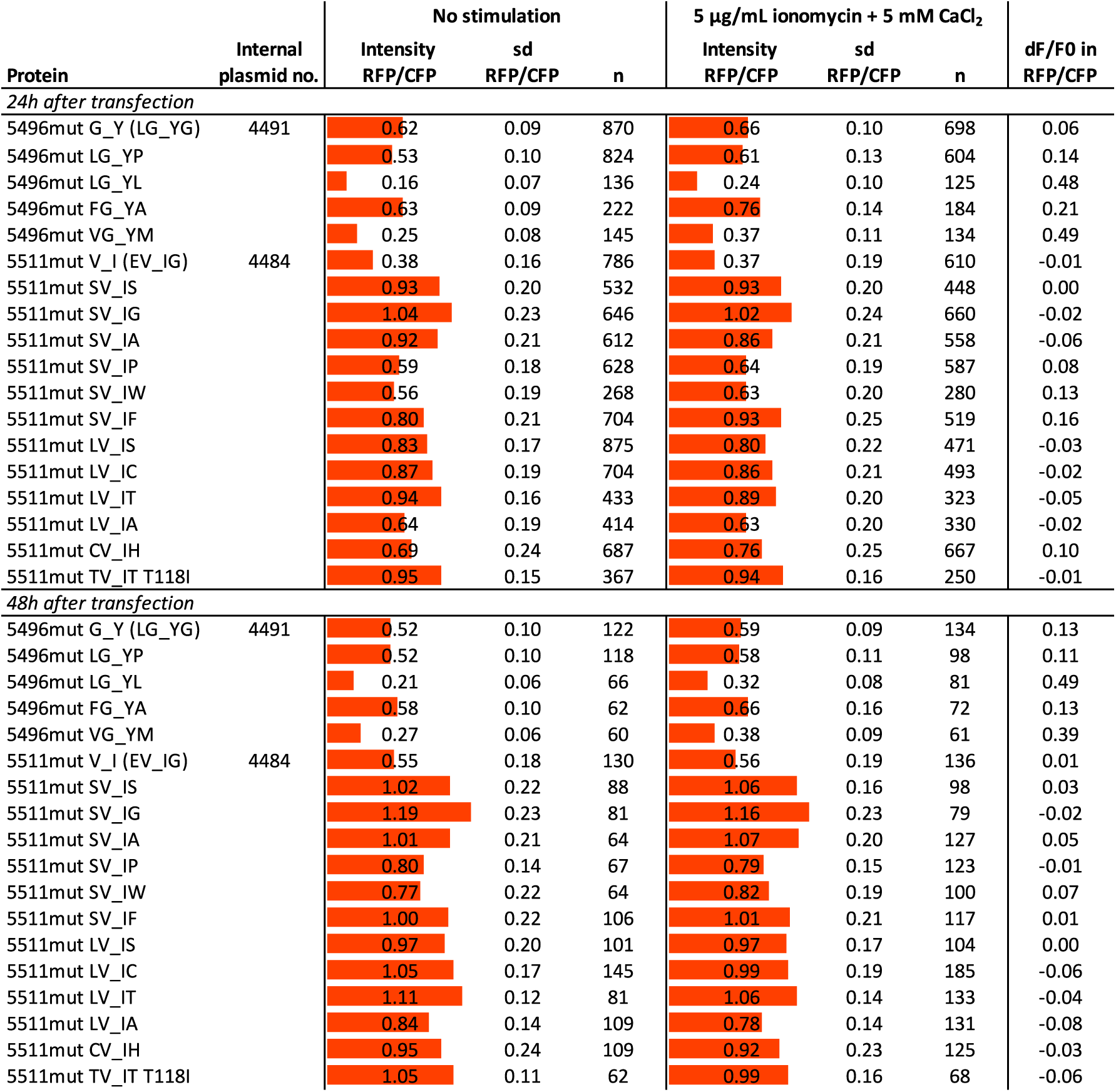
Intensity of 5511mut V_I and 5496mut G_Y mutant sensors in HeLa cells. Red and cyan fluorescence was measured in HeLa cells at 37 °C, at both 24- and 48-hours post-transfection with a pFR plasmid coding a red calcium sensor and mTq2. The ratio RFP/CFP was determined for each cell and normalized to mCherry and no RFP present (only mTq2 expression). Averages of all cells (n) and the standard deviation (sd) are indicated. The measurements were repeated after addition of 5 µg/mL ionomycin and 5 mM CaCl_2_. The change in RFP/CFP ratio divided over the RFP/CFP ratio before addition (d*F*/*F*_0_) was calculated. Mutants are named after their ancestor and mutations, f.e. 5511mut LV_IA stems from variant 5511mut V_I and has mutations E29L and G267A. The original residues at the positions of mutations are indicated at the ancestors 5511mut V_I and 5496mut G_Y.

### Lifetime measurements in HeLa cells

A selection of candidate mScarlet calcium sensors was expressed in HeLa cells, and the fluorescence lifetime before and after addition of ionomycin was recorded (**Figure S6**). The fluorescent phase lifetime of all 5496 variants is between 2.0 and 2.5 ns, both before and after addition. For all 5511 variants, the phase lifetime is between 2.5 and 3.0 ns. In bacterial lysates the best performing sensors showed a lifetime change of over 0.5 ns (calcium-bound minus calcium-free). Unfortunately, only variant 5511 showed a significant but weak positive phase and modulation lifetime change (unstimulated τ_ϕ_ = 2.81 ns [sd = 0.07], τ_M_ = 3.21 ns [sd = 0.12] and high calcium τ_ϕ_ = 2.88 ns [sd = 0.05], τ_M_ = 3.38 ns [sd = 0.16]). For all other variants, the measured changes were absent or non-significant in either the phase or modulation lifetime. The direction of the lifetime changes, if present, is in agreement with measurements in bacterial lysates.

Due to the differences between the measurements in bacterial lysates and HeLa cells, we decided to measure the lifetime change of RCaMP1h and jRCaMP1b in bacterial lysate, using the pFR plasmid (**Table S4**). Both show a much different response in lysates compared to the values in HeLa cells we published before ^21^, namely Δτ_ϕ_ = 0.78 ns, Δτ_M_ = 0.72 ns in lysate versus Δτ_ϕ_ = 1.69 ns, Δτ_M_ = 1.25 ns in HeLa cells for RCaMP1h, and Δτ_ϕ_ = −0.27 ns, Δτ_M_ = −0.12 ns in lysate versus Δτ_ϕ_ = 1.18 ns, Δτ_M_ = 0.67 ns in HeLa cells for jRCaMP1b. As control, we used protein isolate of RCaMPh1h in Tris-HCl buffer, in diluted lysis buffer (with 0.8% DOC, used for screening of 5496mut G_Y and 5511mut V_I mutants) and in pure lysis buffer (2% DOC, used for all other screenings). In Tris buffer, RCaMP1h shows a lifetime change of Δτ_ϕ_ = 1.96 ns and Δτ_M_ = 1.61 ns, but this is reduced in 0.8% DOC (Δτ_ϕ_ and Δτ_M_ = 0.45 ns) and 2% DOC (Δτ_ϕ_ = 0.08 ns, Δτ_M_ = 0.11 ns). In comparison, mScarlet sensor variants 5511, 5496, 5511mut V_I and 5496mut G_Y show almost identical lifetime changes in both concentrations of DOC (**Table S1** for 2% DOC and **Table S3** for 0.8% DOC).

## Discussion

Our aim was to generate a red calcium sensor with a large lifetime contrast based on a mScarlet FP, with improved brightness compared to other red calcium sensors. We were able to generate red fluorescent candidate sensors by circular permutation of different mScarlet FPs, and connecting calmodulin and a calmodulin binding peptide (M13) on respectively the C- and N-terminus. The most successful location for the circular permutation is around the same site (amino acid 145) as all other published red calcium sensors ^6–8,10–15^.

Two designs with mScarlet-I3 showed the most promising results. Number 5511 had the largest fluorescence lifetime change (Δτ_ϕ_ = 0.35 ns, Δτ_M_ = 0.30 ns) between zero calcium and high calcium in bacterial lysate. Number 5496 displayed a much smaller lifetime change (Δτ_ϕ_ = −0.10 ns, Δτ_M_ = −0.03 ns), but visually a good intensity in bacteria. In both designs, we did two rounds of directed random mutagenesis, targeting in total 4 residues. When screening candidate sensors in bacterial lysate after the second round of mutagenesis, we measured fluorescence lifetime changes of Δτ_ϕ_ and Δτ_M_ > 0.5 ns for three variants (5511mut LV_IA, 5511mut LV_IC and 5511mut TV_IA). However, the lifetime response in HeLa cells was only marginal and mostly non-significant. In addition, two variants were identified in the bacterial lysis test with an intensity fold-change of over 20 (5511mut SV_IP and 5511mut SV_IW), but again this was not reflected in HeLa cells.

It was not investigated why the candidate mScarlet sensors perform poorly in mammalian cells compared to bacterial lysate. One possibility is that the *K*_d_ of these sensors is very low. In bacterial lysate the zero-calcium environment puts all sensors in a calcium-free state, while the 100 nM calcium ^25^ in resting HeLa cells might already be enough to saturate (a portion of) the sensors. Sensors with the same calcium binding domains as our mScarlet candidates all have a *K*_d_ in the range of 300–500 nM ^6,21,23^. We find it therefore unlikely that the *K*_d_ of the mScarlet candidates is close to or below 100 nM. However, the difference between 0 and 100 nM calcium between bacterial lysate and HeLa cells will likely have a small influence. A second possibility is that the mild lysis buffer we used for the bacteria is still too harsh for the sensors, changing the conformation slightly and making them more sensitive to the environment. We did measure an influence of DOC on the fluorescence lifetime change of RCaMP1h. However, the contrast for this sensor was reduced with increasing concentrations of DOC. The effect on the mScarlet candidate sensors could be inverse, due to differences between mScarlet and mRuby (in RCaMP1h). The third possibility is that other differences in environment cause a different reaction in the same sensor.

For comparison of the intensity of red sensors, a dedicated plasmid was created that codes for simultaneous expression of equal amounts of mTurquoise2 and a red (candidate) sensor. Remarkably, all candidate sensors from the final rounds of mutagenesis were much brighter than several published sensors that we compared in HeLa cells, namely R-GECO1, K-GECO1, CH-GECO2.1, RCaMP1h and jRCaMP1b. To our knowledge, no quantitative investigation has been conducted on the intensity of red calcium sensors compared to normal FPs like mCherry.

When sensors are published, often a fold-change is reported, determined from protein isolate, for example K-GECO1^14^ (*F*_max_/*F*_0_ = 12) and jRCaMP1b ^11^ (*F*_max_/*F*_0_ = 7.2). In vivo fold-changes were also reported for R-GECO1^6,13^ (in vivo *F*_max_/*F*_0_ 4.9 and 2.5, in vitro *F*_max_/*F*_0_ = 16 and 12.6) and RCaMP1h ^13^ (in vivo *F*_max_/*F*_0_ = 2, in vitro *F*_max_/*F*_0_ = 10.5). We measured fold-changes of 2.7 for R-GECO1 and 2.2 for RCaMP1h and jRCaMP1b after 24 h expression in HeLa cells. Our results again clearly show that intensity changes in vitro do not hold up in mammalian studies. Therefore, it is vital to take this into account when developing a new genetically encoded sensor.

We did notice that the standard deviations of our intensity measurements are high, mainly for the calcium sensors published by others. These sensors have the largest fold-changes, so small differences in the calcium levels between the cells give the largest variation. Possibly the maturation speed of the red sensors is also of influence. If the sensors have not fully matured yet, possibly the readout is different. In our experience, the expression level of a sensor has an influence on the fold-change, with a lower expression level leading to a higher fold-change (not published). For CH-GECO2.1, no intensity fold-change was expected, due to its low *K*_d_ of 6 nM ^15^, below the basal calcium concentration of HeLa cells of 100 nM ^25^. Indeed, we did not measure a fold-change, but still the standard deviation was higher compared to the mScarlet-candidate sensors. It is worthwhile to further improve the intensity comparison, by calculating the fold-change for individual cells instead of on population level, and comparing the fold-change to the expression level.

Although we were not able to develop a successful bright lifetime sensor based on mScarlet, we do still believe in the potential of mScarlet as a scaffold for a calcium sensor. Not only were fluorescent circular permutated variants created of various mScarlet FPs, but also several variants were found that were responsive in bacterial lysate. Especially the brightness of the created candidates is notable. Further improvement is required to reach a sensor that gives these changes in live biological systems, like mammalian cells.

To this end, we propose to switch from bacterial screening to screening in mammalian cells. The fluorescence lifetime will need to be measured before and after raising the calcium level in the same cells. Ideally, a mTurquoise2 fluorescent protein is expressed simultaneously for comparison of the intensity. The pFR plasmid we developed is suitable for the screening process. We anticipate that many new variants need to be created by mutagenesis and compared. So, for high throughput, it is required to fully automate the measurement and stimulation of the cells. As starting point for further improvement, we advise either 5511mut LV_IA, 5511mut LV_IC or 5511mut TV_IA. Their intensity in mammalian cells is bright and they show potential by a high fluorescence phase and modulation lifetime change in bacterial lysate. Alternatively, 5511mut SV_IP or 5511mut SV_IW could be interesting if one desires to develop a sensor with a large intensity change.

## Methods

### General Cloning

We used *E. coli* strain *E. cloni* 5-α (short: *E. cloni*, Lucigen corporation) for all cloning procedures. For DNA assembly, competent *E. cloni* was transformed using a heat shock protocol according to manufacturers’ instructions. For protein expression, *E. cloni* was grown using super optimal broth (SOB, 0.5% (w/v) yeast extract, 2% (w/v) tryptone, 10 mM NaCl, 20 mM MgSO_4_, 2.5 mM KCl) supplemented with 100 µg/mL kanamycin and 0.2% (w/v) rhamnose (SKR). For agar plates, 1.5% (w/v) agar was added. Bacteria were grown overnight at 37 °C. Plasmid DNA was extracted from bacteria using the GeneJET Plasmid Miniprep Kit (Thermo Fisher Scientific) and the obtained concentration was determined by Nanodrop (Life Technologies). DNA fragments were generated by PCR, using Pfu DNA polymerase (Agilent Technologies) unless otherwise indicated. DNA fragments were visualized by gel electrophoresis on a 1% agarose gel, run for 30 min at 80 V. PCR fragments were purified using the GeneJET PCR purification Kit (Thermo Fisher Scientific) and digested with restriction enzymes to generate sticky ends. Restriction enzymes were heat inactivated at 80 °C for 20 min if necessary. Vector fragments were generated by restriction of plasmids and the correct bands were extracted from gel using the GeneJET Gel Extraction Kit (Thermo Fisher Scientific). DNA fragments were ligated using T4 DNA ligase (Thermo Fisher Scientific), per the manufacturers’ protocol. For targeted mutagenesis, we used the protocol as described before ^26^. In short, primers were designed with an annealing region of 15 bp both left and right of the desired mutation(s). The full plasmid was amplified by PCR and the DNA was digested with DpnI to remove template DNA. *E. cloni* was transformed with the digested PCR mix and plasmids were isolated from individual clones. Correct construction of plasmids was verified by control digestion and sequencing (Macrogen Europe). All primers (**Table S5**) were ordered from Integrated DNA Technologies.

### Construction of sensors with mScarlet-I/mScarlet-A220

Circular permutated variants of mScarlet-I (cp-ScI) and a mutant of mScarlet (G220A, cpScA220) were constructed by PCR amplification from a tandem construct containing two mScarlet-I (td-ScI) or mScarlet-A220 (td-ScA220) proteins connected by a flexible GGSGG-linker (primers 1–25). Primers contain the sequence that connects the circular permutated FP to the M13 or the CaM on the N- or C-terminal side, respectively, by linker sequences. The linking sequences were based on the linking sequences in R-GECO1 and CH-GECO (primers 1–17 and 18–25, respectively). Primers that annealed on various positions on mScarlet-I or mScarlet-A220 and with different length were used to create different variants. The cpmTurquoise2 in pFHL-Tq-Ca-FLITS (Addgene plasmid #129628) was replaced with different cpScI or cpScA220 variants by digestion of the vector and the PCR fragments with SacI and MluI, followed by ligation. Correct construction was verified by sequencing (primers seq1 and seq2, see **Table S5**).

A few directed mutations were added on some of the mutants, to better mimic the R-GECO1 sensor. These were: T148S (primer 11), M164K and M164R (primers 26–27). The M164K/R mutations were created by directed mutagenesis), see “General cloning” for details. The fluorescent protein in K-GECO1 (Addgene plasmid #105864, a gift from Robert Campbell ^14^) was replaced with a cpScI and cpScA220 by in vitro assembly ^27,28^. An insert and backbone fragment were created by PCR in a single PCR tube containing all four primers (primers 28–31) and both templates (pFHL-Tq-Ca-FLITS and a plasmid containing td-ScI or td-ScA220). The PCR mix was digested with DpnI to remove template DNA and 50 µL heat shock competent *E. cloni* was transformed with 6 µL of this mix. Correct construction of the plasmids was evaluated by control digestion with SacII and PvuII and MfeI and EagI, and sequencing (primers seq1-seq5). A non-responsive variant was created as a control with a non-circular permutated mSc-A220 FP (primers 32–33) in between the M13-peptide and the CaM domain, named cp1ScA220-GECO. For a full list of all sensors created, see **Table S6**.

### Construction of sensors with mScarlet-I3/mScarlet3

After trial and error, the cpScA220 in the best performing candidate sensor so far (number 5309) was replaced by circular permutated mScarlet-I3 (cpScI3) and mScarlet3 (cpSc3), using the same position of permutation and the same linking sequences. In addition, 10 other almost identical variants were created, with single amino acid insertions and deletions. The in total 12 new variants were created as described above using primers 4, 6, 7, 9, 18, 19, 34–39 (**Table S5**). The sequences were determined by sequencing using primers seq1 and seq2. To create a library of sensors with different linkers, a similar approach was taken, but now a library of cpScI3 fragments was generated by PCR using a mix of primers (primers 1–11, 18, 19, 34–39). Based on the results of screening the library, we rationally designed and created three more variants (primers 1, 19, 38 and 40). Interesting sensor variants 5496 and 5511 were subjected to targeted random mutagenesis on two positions in the linking sequences, generating 400 possible variants of each sensor. The same cloning approach was taken, now using primers containing altered or degenerated codons (primer 41 and 42 for mutagenesis of variant 5511, and primers 42 and 43 for variant 5496). On four of the resulting variants, we did a directed mutagenesis of R198I (mScarlet numbering), using primers 44 and 45, see “General cloning” for details. The best candidates were selected and subjected to targeted random mutagenesis on two other positions. Again, we used degenerate primers, generating 400 possible variants (primers 46 and 47 for mutagenesis of 5496mut G_Y, and primers 48 and 49 for 5511mut V_I). For a full list of all variants, see **Table S6**. Sequencing of all ScI3 and Sc3 sensor variants was done using primers seq1 and/or seq2.

### Construction of a dual expression vector for ratiometric measurements

The SacI restriction site was removed from mTq2 in an “empty” pDress plasmid ^22^, containing mTq2 linked to an anti-FRET linker and a P2A site (Addgene plasmid #130509, but with the mScarlet protein between the AgeI and BsrGI sites replaced with the DNA sequence GCCTCAACATGA). This was achieved by directed mutagenesis using primers 50–51, see “General cloning” for details. After mutagenesis, correct removal of the SacI site and the absence of unwanted spontaneous mutations was verified by control digestion with SacI and NheI followed by sequencing (primers 5–7). This yielded pDress_mTq2_antiFRET_P2A_linker-noSacI (internal plasmid number 5516). An insert fragment was amplified from the new plasmid by PCR (primers 52–53), containing the last part of the rhamnose promotor, the ribosome biding site, the adapted kozak sequence, the 6xHis-tag, the thrombin recognition site, the mTq2 fluorescent protein, the anti-FRET linker and the P2A sequence ^22^. A PCR (primers 54–55) with Phusion polymerase (Thermo Fisher Scientific) was performed on another dual expression plasmid pFHL (Addgene plasmid #129628) to generate a backbone fragment containing a calcium sensor, the backbone of the pFHL plasmid, the CMV promotor and the first part of the rhamnose promotor. Both insert and backbone fragment were digested with BamHI and EcoRI to create overhanging ends. The fragments were ligated together and *E. cloni* was transformed with the DNA. Correct construction of the new pFR (Franka Ratio) plasmid was verified by control digestion with NheI and BsrGI and sequencing (primers seq2, seq7-seq8).

### Placement of red calcium sensors in the ratioplasmid

R-GECO1 (Addgene plasmid #32465, a gift from Robert Campbell ^6^), CH-GECO2.1 (Addgene plasmid #52099 a gift from Robert Campbell ^15^), K-GECO1 (Addgene plasmid #105864, a gift from Robert Campbell ^14^), jRCaMP1b (Addgene plasmid #63136, a gift from Douglas Kim & GENIE Project ^11^) and RCaMP1h (Addgene plasmid #42874, a gift from Loren Looger ^13^) were placed in the pFR plasmid. First, insert fragments were generated by PCR, using primers 56–57 for K-GECO1 and primers 57–58 for all others (**Table S5**). Next, both inserts and a pFR were digested with BamHI and HindIII, and the inserts were ligated into the backbone. Finally, correct insertion was verified by sequencing (primers seq7-seq9). In a similar manner mCherry (pmCherry-C1) was placed in pFR, using primers 59–60 for construction.

### Testing pFR for use in bacteria and mammalian cells

The performance of pFR was tested in three situations: i) in bacteria on agar plates, ii) in bacterial lysate and iii) in mammalian cells.

1. Bacteria expressing mTq2 (internal plasmid number #5516) or mTq2-cp1mScA220-GECO (internal plasmid number #5521) were grown on agar plates containing kanamycin and rhamnose (for expression). The lifetime of the CFP was determined using a home-build Zeiss setup controlled by Matlab 6.1 software, composed of an Axiovert 200M inverted fluorescence microscope (Zeiss) with a II18MD modulated image intensifier (Lambert Instruments) coupled to a CoolSNAP HQ CCD camera (Roper Scientific) and two computer-controlled HF-frequency synthesizers (SML 01, Rohde & Schwartz), one driving the intensifier and the other driving a 440 nm modulated laser diode (PicoQuant, LDH-M-C-440) through a DL-300 driver unit ^29^. The excitation light is modulated at 75.1 MHz and reflected by a 455 nm dichroic mirror onto the sample. Emission is filtered with a 480/40 nm band-pass emission filter. A F100 lens (Spindler & Hoyer) was used. Data were analyzed as described before ^30^ using an imageJ macro.
2. From an overnight 2.5 mL liquid culture of the same bacteria, a crude lysate was extracted similar as described before ^23^. The bacteria were collected by centrifugation for 1 min at 13.2 ×g and 800 µL of lysis buffer (2% Deoxycholic acid in 50 mM Tris-HCl pH 8.0) was added. The bacteria were resuspended and left to incubate for 15 min, while intermittently shaking. The suspension was centrifuged for 10 min at 13.2 ×g and the crude lysate was collected. The lysate was diluted 100–500× in lysis buffer until a faint yellow color was visible. Either 0.10 mM CaCl_2_ or 9.5 mM EDTA was added and the lifetime of the CFP in the lysate was measured as described under “Lifetime imaging.” The lifetime measurements were repeated after a photobleaching pulse of 1 min from a HG-light source (Intensilight C-HGFIE, Nikon) at maximum intensity with a 609/54 nm excitation filter and a 561 nm dichroic mirror (Semrock).
3. Mammalian cells were grown and transfected as described under “HeLa cell culture and transfection.” The fluorescence lifetime of the CFP was measured as described under “Lifetime imaging.” The measurements were repeated after a photobleaching pulse as described above. To other samples, first ionomycin (5 µg/mL, I-6800, LClaboratories) and extra calcium (5 mM) were added, followed by the same series of measurements.

### Bacterial screening

*E. cloni* bacteria were used for screening, both after generation of a pool of mScarlet-I3 sensor variants with different linkers and after targeted random mutagenesis. First, bacteria expressing sensor variants from a pFHL or pFR plasmid were grown SKR-agar plates. Colonies visually showing a (bright) red fluorescence under a binocular (excitation filter 580/20 nm, emission filter 645/75 nm) under a stereomicroscope (MZFLIII, Leica) were selected for a bacterial lysis test. Here, bacteria were grown overnight in 1.5 mL SKR in a deep wells plate (732-2893, VWR) covered with an adhesive seal (AB-0718, Thermo Fisher Scientific) at 37 °C while shaking at 280 rpm. The second day, 1 µL of culture was spotted on a SKR plate for storage, and the remainder of the bacteria were harvested by 15 min centrifugation at 2683 ×g in a swing-out centrifuge (5810 R, Eppendorf). Supernatant was removed and 200 µL of lysis buffer (2% Deoxycholic acid in 50 mM Tris-HCl pH 8.0) was added. Bacteria were brought in suspension using a vortex and incubated for 15 minutes at RT while intermittently shaking. Cell debris was removed by 40 min centrifugation at 2683 ×g in a swing-out centrifuge. The bacterial lysate was collected and stored overnight at 4 °C.

The third day, bacterial lysates were diluted 2.5× in 50 mM Tris-HCl pH 8.0 (for screening of 5511mut V_I and 5496mut G_Y) or lysis buffer (all other lysates) to a final volume of 200 µL in 96-well plates with black walls and a glass bottom (89626, Ibidi), unless otherwise specified. Red fluorescence and, if applicable, cyan fluorescence were measured using a FL600 microplate fluorescence reader controlled by KC4™ software (Bio-Tek) with 430/25 or 555/25 nm excitation and 485/40 or 620/40 nm emission for, respectively, CFP or RFP, and averaging each well 10×. The value of a background well (the same buffer as used for dilution) was subtracted and the RFP intensity divided over CFP (if applicable). The same plate was used for determination of the fluorescence lifetime of the lysates as described under “Lifetime imaging.” Intensity measurements and lifetime measurements were repeated after addition of 0.1 mM CaCl_2_ and after a second addition of 9.5 mM EDTA. Intensity fold-change was calculated as the RFP/CFP intensity with calcium divided over RFP/CFP without calcium.

### HeLa cell culture and transfection

HeLa cells (CCL-2) acquired from the American Tissue Culture Collection were maintained in full medium, Dulbecco’s modified Eagle medium + GlutaMAX (61965, Gibco) supplemented with 10% fetal bovine serum (FBS, 10270, Gibco), under 7% humidified CO_2_ atmosphere at 37 °C. Cells were washed with Hank’s buffered salt solution (HBSS, 14175, Gibco) and trypsinized (25300, Gibco) for passaging. No antibiotics were used. HeLa cells were grown in 24-wells plates with a glass bottom for imaging (Thermo Fisher Scientific). Transfection mixture was prepared in Opti-MEM (31985047, Thermo Fisher Scientific) with 2 µg Polyethylenimine in water (PEI, pH 7.3, 23966, Polysciences), 50 ng plasmid DNA and 150 ng mock-DNA (empty plasmid), and incubated for 20 min before addition to the cells. Cells were imaged 1- or 2-days post-transfection. Prior to lifetime measurements, the medium was replaced with microscopy medium (137 mM NaCl, 5.4 mM KCl, 1.8 mM CaCl_2_, 0.8 mM MgSO_4_, 20 mM D-Glucose, 20 mM HEPES pH 7.4) and cells were incubated for 20 min.

### Lifetime imaging

For lifetime measurements, we used a Lambert Instruments FLIM Attachment (LIFA) setup, composed of an Eclipse Ti microscope (Nikon) with a Lambert Instruments Multi-LED for excitation, a LI2CAM camera, a LIFA signal generator (all Lambert Instruments) to synchronize the light source and the camera, and was controlled by the LI-FLIM software (version 1.2.13). For RFP measurements, a 532 nm light emitting diode (LED) was used, combined with a 534/20 nm excitation filter, a 561 nm dichroic mirror and a 609/54 nm band-pass filter. For CFP measurements, a 446 nm LED was used, combined with a 448/20 nm excitation filter, a 442 nm dichroic mirror and a 482/25 nm band-pass filter (all filters from Semrock). Alexa488, mScarlet or EB was used as a reference to calibrate the instrumentation, with a known mono-exponential lifetime of 4.05 ns ^31–33^, 3.86 ns ^16^ or 0.086 ns ^34–36^, respectively.

Fluorescence lifetime in HeLa cells was recorded at 37 °C before and after addition of a mix of ionomycin (5 µg/mL, I-6800, LClaboratories) and CaCl_2_ (5 mM). Cells were imaged using a ×40 (Plan Apo, NA 0.95 air) objective and collecting 12 phase images. Fluorescence lifetime in bacterial lysates were recorded at RT using a ×20 (Plan Apo, NA 0.95 air) objective, collecting 12 phase images and averaging 3×. When measuring lysates in the previously described 96-wells plate, the LIFA software was controlled by a Matlab script ^22^ that automatically moves to the position of each of the wells, adjusts the exposure time based on the intensity to a maximum of 500 ms, collects the lifetime stack and saves the data.

Recorded sample stacks and a reference stack were converted into lifetime images by an ImageJ macro ^22,37^. When the intensity of fluorescence in HeLa cells was very weak, a background correction was performed on the lifetime stacks, using a manually indicated background region using Matlab (R2015a). For bacterial lysates, the average phase and modulation lifetime of the full view were extracted using the imageJ macro. For cells, the lifetime of individual cells with was collected. This was done in a semi-automatic manner, using first an ImageJ macro that masks the image based on intensity, followed by a macro that guides the measurements of the cells and the saving of the ROIs.

### Ratiometric imaging

HeLa cells co-expressing a red sensor (or red FP) and mTq2 from a single plasmid (pFR) were imaged 24 h or 48 h after transfection, at 37 °C, using an Eclipse Ti microscope (Nikon) equipped with a Spectra X Light Engine (Lumencor) for excitation and an Orca flash 4.0 camera (Hamamatsu). Cells were imaged using a Plan Apo ×10 NA 0.45 air objective with binning 2×2 and only imaging the middle of the camera, or by using a Plan Apo ×20 NA 0.75 air objective and binning 4×4. For imaging of RFP, a 575 nm LED, a 575/25 nm excitation filter, and a quad band cube (MXU 71640, Nikon) were used. For CFP, a 440 nm LED, a 440/20 nm excitation filter, and triple quad cube (MXU 74157, NIKON) were used. Channels were imaged sequentially. Fluorescence was recorded before and directly after addition of a mix of ionomycin (5 µg/mL, I-6800, LClaboratories) and calcium (5 mM).

Background was subtracted from the images and the average red and cyan fluorescence intensity of individual cells was measured using ImageJ (version 1.52k). The ratio RFP/CFP was calculated for each cell. As a controls, we used a sample with only mTq2 (ratio RFP/CFP = 0) and a sample with the dummy sensor (cp1mScA220-GECO) and mTq2. The ratio of all cells was corrected to the controls and normalized to mCherry. The experiment was performed twice per construct and timepoint.

### Protein isolation of RCaMP1h

His-tagged RCaMP1h was isolated from bacterial culture essentially as described before, using Ni^2+^ loaded His-Bind resin ^26^. In the final step, the isolated protein was overnight dialyzed in 10 mM Tris-HCl pH 8.0. No further purification was performed. Proteins were snap frozen in liquid nitrogen for long term storage at −80 °C.

## Acknowledgements

F.H.L. was supported by a NWO Chemical Sciences ECHO grant (711.017.003).

## Author contributions

F.H.L., J.G. and T.W.J.G. conceptualized the project and designed the experiments. F.H.L. performed all experiments, analyzed and interpreted the results, and wrote the manuscript.

## Data availability

Several plasmids are deposited for distribution through Addgene (www.addgene.org). The plasmids and corresponding addgene numbers are: pFR-mTq2-antiFRET-P2A-mCherry: #191466, pFR-mTq2-antiFRET-P2A-RCaMP1h: #191467, pFR-mTq2-antiFRET-P2A-jRCaMP1b: #191468, pFR-mTq2-antiFRET-P2A-K-GECO1: #191469, pFR-mTq2-antiFRET-P2A-CH-GECO2.1: #191470, pFR-mTq2-antiFRET-P2A-R-GECO1: #191471, pDress-mTq2-antiFRET-P2A2-linker_noSacI: #191472.

## Notes

### Competing Interest Statement

The authors have declared no competing interest.

## References

[1] E.C. Greenwald, M. Sohum, and J. Zhang. Genetically Encoded Fluorescent Biosensors Illuminate the Spatiotemporal Regulation of Signaling Networks. Chemical Reviews, 118:11707–11794, 2018.

[2] G.S. Baird, D.A. Zacharias, and R.Y. Tsien. Circular permutation and receptor insertion within green fluorescent proteins. Proceedings of the National Academy of Sciences, 96:11241–11246, 1999.

[3] O. Griesbeck, G.S. Baird, R.E. Campbell, D.A. Zacharias, and R.Y. Tsien. Reducing the environmental sensitivity of yellow fluorescent protein. Mechanism and applications. Journal of Biological Chemistry, 276:29188–29194, 2001.

[4] J. Nakai, M. Ohkura, and K. Imoto. A high signal-to-noise Ca2+ probe composed of a single green fluorescent protein. Nature Biotechnology, 19:137–141, 2001.

[5] T. Nagai, A. Sawano, E.S. Park, and A. Miyawaki. Circularly permuted green fluorescent proteins engineered to sense Ca2+. Proceedings of the National Academy of Sciences, 98:3197–3202, 2001.

[6] Y. Zhao, S. Araki, J. Wu, T. Teramoto, Y.F. Chang, M. Nakano, A.S. Abdelfattah, M. Fujiwara, T. Ishihara, T. Nagai, and R.E. Campbell. An Expanded Palette of Genetically Encoded Ca2+ indicators. Science, 333:1888–1891, 2011.

[7] M. Ohkura, T. Sasaki, C. Kobayashi, Y. Ikegaya, and J. Nakai. An improved genetically encoded red fluorescent Ca2+ indicator for detecting optically evoked action potentials. PLoS ONE, 7, 2012.

[8] J. Wu, L. Liu, T. Matsuda, Y. Zhao, A. Rebane, M. Drobizhev, Y.F. Chang, S. Araki, Y. Arai, K. March, T.E. Hughes, K. Sagou, T. Miyata, T. Nagai, W.H. Li, and R.E. Campbell. Improved orange and red Ca2+indicators and photophysical considerations for optogenetic applications. ACS Chemical Neuroscience, 4:963–972, 2013.

[9] J. Suzuki, K. Kanemaru, K. Ishii, M. Ohkura, Y. Okubo, and M. Iino. Imaging intraorganellar Ca2+at subcellular resolution using CEPIA. Nature Communications, 5:1–13, 2014.

[10] M. Inoue, A. Takeuchi, S.I. Horigane, M. Ohkura, K. Gengyo-Ando, H. Fujii, S. Kamijo, S. Takemoto-Kimura, M. Kano, J. Nakai, K. Kitamura, and H. Bito. Rational design of a high-affinity, fast, red calcium indicator R-CaMP2. Nature Methods, 12:64–70, 2015.

[11] H. Dana, B. Mohar, Y. Sun, S. Narayan, A. Gordus, J.P. Hasseman, G. Tsegaye, G.T. Holt, A. Hu, D. Walpita, R. Patel, J.J. Macklin, C.I. Bargmann, M.B. Ahrens, E.R. Schreiter, V. Jayaraman, L.L. Looger, K. Svoboda, and D.S. Kim. Sensitive red protein calcium indicators for imaging neural activity. eLife, 5:1–24, 2016.

[12] O.M. Subach, N.V. Barykina, E.S. Chefanova, A.V. Vlaskina, V.P. Sotskov, O.I. Ivashkina, K.V. Anokhin, and F.V. Subach. Frcamp, a red fluorescent genetically encoded calcium indicator based on calmodulin from schizosaccharomyces pombe fungus. International Journal of Molecular Sciences, 22, 2021.

[13] J. Akerboom, N. Carreras Calderón, L. Tian, S. Wabnig, M. Prigge, J. Tolö, A. Gordus, M.B. Orger, K.E. Severi, J.J. Macklin, R. Patel, S.R. Pulver, T.J. Wardill, E. Fischer, C. Schüler, T.W. Chen, K.S. Sarkisyan, J.S. Marvin, C.I. Bargmann, D.S. Kim, S. Kügler, L. Lagnado, P. Hegemann, A. Gottschalk, E.R. Schreiter, and L.L. Looger. Genetically encoded calcium indicators for multi-color neural activity imaging and combination with optogenetics. Frontiers in Molecular Neuroscience, 6:1–29, 2013.

[14] Y. Shen, H. Dana, A.S. Abdelfattah, R. Patel, J. Shea, R.S. Molina, B. Rawal, V. Rancic, Y.F. Chang, L. Wu, Y. Chen, Y. Qian, M.D. Wiens, N. Hambleton, K. Ballanyi, T.E. Hughes, M. Drobizhev, D.S. Kim, M. Koyama, E.R. Schreiter, and R.E. Campbell. A genetically encoded Ca2+indicator based on circularly permutated sea anemone red fluorescent protein eqFP578. BMC Biology, 16:1–16, 2018.

[15] H.J. Carlson and R.E. Campbell. Circular permutated red fluorescent proteins and calcium ion indicators based on mCherry. *Protein Engineering*, Design and Selection, 26:763–772, 2013.

[16] D.S. Bindels, L. Haarbosch, L. Van Weeren, M. Postma, K.E. Wiese, M. Mastop, S. Aumonier, G. Gotthard, A. Royant, M.A. Hink, and T.W. Gadella. MScarlet: A bright monomeric red fluorescent protein for cellular imaging. Nature Methods, 14:53–56, 2016.

[17] T.W. Gadella, L. van Weeren, J. Stouthamer, M.A. Hink, A.H. Wolters, B.N. Giepmans, S. Aumonier, J. Dupuy, and A. Royant. mScarlet3: a brilliant and fast-maturing red fluorescent protein. Nature Methods, 20:541–545, 2023.

[18] A. Liu, X. Huang, W. He, F. Xue, Y. Yang, J. Liu, L. Chen, L. Yuan, and P. Xu. pHmScarlet is a pH-sensitive red fluorescent protein to monitor exocytosis docking and fusion steps. Nature Communications, 12:1–12, 2021.

[19] C. Beck and Y. Gong. A high-speed, bright, red fluorescent voltage sensor to detect neural activity. Scientific Reports, 9:1–12, 2019.

[20] C.M. Díaz-García, R. Mongeon, C. Lahmann, D. Koveal, H. Zucker, and G. Yellen. Neuronal Stimulation Triggers Neuronal Glycolysis and Not Lactate Uptake. Cell Metabolism, 26:361–374, 2017.

[21] F.H. van der Linden, E.K. Mahlandt, J.J. Arts, J. Beumer, J. Puschhof, S.M. de Man, A.O. Chertkova, B. Ponsioen, H. Clevers, J.D. van Buul, M. Postma, T.W. Gadella, and J. Goedhart. A turquoise fluorescence lifetime-based biosensor for quantitative imaging of intracellular calcium. Nature Communications, 12:1–13, 2021.

[22] D.S. Bindels, M. Postma, L. Haarbosch, L. van Weeren, and T.W. Gadella. Multiparameter screening method for developing optimized red-fluorescent proteins. Nature Protocols, 15:450–478, 2020.

[23] F.H. van der Linden, S.C. Thornquist, R.M. ter Beek, J.Y. Huijts, M.A. Hink, T.W. Gadella, G. Maimon, and J. Goedhart. A green lifetime biosensor for calcium that remains bright over its full dynamic range. bioRxiv, 2024.

[24] J. Goedhart, D. Von Stetten, M. Noirclerc-Savoye, M. Lelimousin, L. Joosen, M.A. Hink, L. Van Weeren, T.W. Gadella, and A. Royant. Structure-guided evolution of cyan fluorescent proteins towards a quantum yield of 93%. Nature Communications, 3, 2012.

[25] M.J. Berridge, P. Lipp, and M.D. Bootman. The Versatility and Universality of Calcium Signalling. Nature Reviews, 1:11–21, 2000.

[26] D.S. Bindels, J. Goedhart, M.A. Hink, L.V. Weeren, L. Joosen, and T.W.J. Gadella. Chapter 16: Optimization of Fluorescent Proteins, volume 1076. Methods in Molecular Biology, 2014.

[27] J. García-Nafría, J.F. Watson, and I.H. Greger. IVA cloning: A single-tube universal cloning system exploiting bacterial In Vivo Assembly. Scientific Reports, 6:27459, 2016.

[28] M. Kostylev, A.E. Otwell, R.E. Richardson, and Y. Suzuki. Cloning should be simple: Escherichia coli DH5á-mediated assembly of multiple DNA fragments with short end homologies. PLoS ONE, 10:1–15, 2015.

[29] E.B. van Munster and T.W. Gadella. фFLIM: A new method to avoid aliasing in frequency-domain fluorescence lifetime imaging microscopy. Journal of Microscopy, 213:29–38, 2004.

[30] E.M. Merzlyak, J. Goedhart, D. Shcherbo, M.E. Bulina, A.S. Shcheglov, A.F. Fradkov, A. Gaintzeva, K.A. Lukyanov, S. Lukyanov, T.W. Gadella, and D.M. Chudakov. Bright monomeric red fluorescent protein with an extended fluorescence lifetime. Nature Methods, 4:555–557, 2007.

[31] E. Rusinova, V. Tretyachenko-Ladokhina, O.E. Vele, D.F. Senear, and J.B. Alexander Ross. Alexa and Oregon Green dyes as fluorescence anisotropy probes for measuring protein-protein and protein-nucleic acid interactions. Analytical Biochemistry, 308:18–25, 2002.

[32] Y. Ni and E. Terpetschnig. Time-Domain Lifetime Measurements on ChronosBH, 2018.

[33] K. Zheng, T.P. Jensen, and D.A. Rusakov. Monitoring intracellular nanomolar calcium using fluorescence lifetime imaging. Nature Protocols, 13:581–597, 2018.

[34] N. Boens, Q. Wenwu, N. Basarić, J. Hofkens, M. Ameloot, J. Pouget, J.P. Lefévre, B. Valeur, E. Gratton, M. VandeVen, N.D. Silva, Y. Engelborghs, K. Willaert, A. Sillen, G. Bumbles, D. Phillips, A.J.W.G. Visser, A. van Hoek, J.R. Lakowicz, H. Malak, I. Gryczynski, A.G. Szabo, D.T. Krajcarski, N. Tamai, and A. Miura. Fluorescence Lifetime Standards for Time and Frequency Domain Fluorescence Spectroscopy. Analytical Chemistry, 79:2137–2149, 2007.

[35] P.I. Bastiaens, W.F. Wolkers, A.J. Visser, A. Van Hoek, and J.C. Brochon. Comparison of the Dynamical Structures of Lipoamide Dehydrogenase and Glutathione Reductase by Time-Resolved Polarized Flavin Fluorescence. Biochemistry, 31:7050–7060, 1992.

[36] E.B. van Munster and T.W. Gadella. Suppression of photobleaching-induced artifacts in frequency-domain FLIM by permutation of the recording order. Cytometry, 58A:185–194, 2004.

[37] T.W.J. Gadella, R.M. Clegg, and T.M. Jovint. Fluorescence lifetime imaging microscopy: pixel-by-pixel analysis of phase-modulation data. Bioimaging, 2:139–159, 1994.

