## Supplemental data for "Exploration of mScarlet for development of a red lifetime sensor for calcium imaging"

**This supplement contains**

- 6 Supplemental Figures
- 6 Supplemental Tables

**Alignment all mScarlet-I and mScarlet-A220 variants**

```
5343, 5344      -----MVDSSRRKWNKAGHAVRAIGRLSS----KYNTERLYPEDGVLKGDIKMALRLKDGGRYLADFKTTYKAKKP---VQMPGAYNVDRK
K-GECO1         MASRMGSVKLIPSLTTVILVKSMLRKRSFGNPFKYNTETLYPADGGLEGACDMALKLVGGGHLNCSLETTYRSKKPATNLKMPGVYNVDHR
R-GECO1         -----MVDSSRRKWNKAGHAVRAIGRLSSPVV----SERMYPEDGALKSEIKKGLRLKDGGHYAAEVKTTYKAKKP---VQLPGAYIVDIK
CH-GECO2.1      -----MVDSSRRKWNKAGHAVRAIGRLSSLES-LLSTERLYPEDGALKGEIKQRLKLKDGGHYAAEVKTTYKAKKP---VQLPDAYIVDIK
5301, 1629      -----MVDSSRRKWNKAGHAVRAIGRLSSPVVWEASTERLYPEDGVLKGDIKMALRLKDGGRYLADFKTTYKAKKP---VQMPGAYNVDRK
5302, 1630      -----MVDSSRRKWNKAGHAVRAIGRLSSPVV-EASTERLYPEDGVLKGDIKMALRLKDGGRYLADFKTTYKAKKP---VQMPGAYNVDRK
5303, 1631      -----MVDSSRRKWNKAGHAVRAIGRLSSPVV--ASTERLYPEDGVLKGDIKMALRLKDGGRYLADFKTTYKAKKP---VQMPGAYNVDRK
5304, 1632      -----MVDSSRRKWNKAGHAVRAIGRLSSPVV---STERLYPEDGVLKGDIKMALRLKDGGRYLADFKTTYKAKKP---VQMPGAYNVDRK
5305, 5300      -----MVDSSRRKWNKAGHAVRAIGRLSSPVV----TERLYPEDGVLKGDIKMALRLKDGGRYLADFKTTYKAKKP---VQMPGAYNVDRK
5309            -----MVDSSRRKWNKAGHAVRAIGRLSSLES-LLSTERLYPEDGVLKGDIKMALRLKDGGRYLADFKTTYKAKKP---VQMPGAYNVDRK
5339, 5340      -----MVDSSRRKWNKAGHAVRAIGRLSSPVV----TERLYPEDGVLKGDIKMALRLKDGGRYLADFKTTYKAKKP---VQMPGAYNVDRK
5341, 5342      -----MVDSSRRKWNKAGHAVRAIGRLSSPVV----SERLYPEDGVLKGDIKMALRLKDGGRYLADFKTTYKAKKP---VQMPGAYNVDRK
5306            -----MVDSSRRKWNKAGHAVRAIGRLSSPVV------------------IKMALRLKDGGRYLADFKTTYKAKKP---VQMPGAYNVDRK
5307            -----MVDSSRRKWNKAGHAVRAIGRLSSPVV----------------------LRLKDGGRYLADFKTTYKAKKP---VQMPGAYNVDRK
5308            -----MVDSSRRKWNKAGHAVRAIGRLSSPVV---------------------------------------------------------RK
5310            -----MVDSSRRKWNKAGHAVRAIGRLSSLES-LL---------------IKMALRLKDGGRYLADFKTTYKAKKP---VQMPGAYNVDRK
5311            -----MVDSSRRKWNKAGHAVRAIGRLSSLES-LL-------------------LRLKDGGRYLADFKTTYKAKKP---VQMPGAYNVDRK
5312            -----MVDSSRRKWNKAGHAVRAIGRLSSLES-LL------------------------------------------------------RK
Aa in mScarlet                                     147            162 164                                   198

K-GECO1         LERIKEADDETYVELHEVAVARYVGLGGG----GGTGG-SVSE-------LIKENMPMKLYMEGTVNNHHFKCTSEGEGKPYEGTQTMRIK
R-GECO1         LDIVSHNEDYTIVEQCERAEGRHSTGGMDELYKGGTGGSLVSKGEEDNMAIIKEFMRFKVHMEGSVNGHEFEIEGEGEGRPYEAFQTAKLK
CH-GECO2.1      LDIVSHNEDYTIVEQYERAEGRHSTGGMDELYKGGTGGSMVSKGVEDNMAFIKEFMRFKVHMEGSVNGHEFEIEGEGEGRPYEGTQTAKLK
1629-1632,5300,5339,5341,5343
                LDITSHNEDYTVVEQYERSEGRHSTGGMDELYQGGSGG-MVSKGE----AVIKEFMRFKVHMEGSMNGHEFEIEGEGEGRPYEGTQTAKLK
5301-5305,5309,5340,5342,5344,5306,5307,5308,5310,5311,5312
                LDITSHNEDYTVVEQYERSEARHSTGGMDELYQGGSGG-MVSKGE----AVIKEFMRFKVHMEGSMNGHEFEIEGEGEGRPYEGTQTAKLK
Aa in mScarlet                      220

K-GECO1         VVEGGPLPFAFDILATSFMYGSRTFIKHPPGIPDFFKQSFPEGFTWERVTTYEDGGVLTATQDTSLQDGCLIYNVKVRGMNFPANGPVMQK
R-GECO1         VTKGGPLPFAWDILSPQFMYGSKAYIKHPADIPDYFKLSFPEGFRWERVMNFEDGGIIHVNQDSSLQDGVFIYKVKLRGTNFPPDGPVMQK
CH-GECO2.1      VTKGGPLPFAWDILSPQFMYGSKAYIKHPADIPDYWKLSFPEGFKWERVMNFEDGGVVTVTQDSSLQDGEFIYKVKLRGTNFPSNGPVMQK
1629-1632,5300,5339,5341,5343
                VTKGGPLPFSWDILSPQFMYGSRAFIKHPADIPDYYKQSFPEGFKWERVMNFEDGGAVTVTQDTSLEDGTLIYKVKLRGTNFPPDGPVMQK
5301-5305,5309,5340,5342,5344,5306,5307,5308,5310,5311,5312
                VTKGGPLPFSWDILSPQFMYGSRAFTKHPADIPDYYKQSFPEGFKWERVMNFEDGGAVTVTQDTSLEDGTLIYKVKLRGTNFPPDGPVMQK
Aa in mScarlet                           74

5343, 5344      KTMGWEA------------------------------------------------------SNGQLTEEQIAEFKEAFSLFDKDGDGTITT
K-GECO1         KTLGWEA------------------------------------------------------SNGQLTEEQIAEFKEAFSLFDKDGDGTITT
R-GECO1         KTMGWEA------------------------------------------------------TRDQLTEEQIAEFKEAFSLFDKDGDGTITT
CH-GECO2.1      KTMGW-----------------------------------------------------LGGTRDQLTEEQIAEFKEAFSLFDKGGAGTITT
5301, 1629      KTMG---------------------------------------------------------TRDQLTEEQIAEFKEAFSLFDKDGDGTITT
5302, 1630      KTMGW--------------------------------------------------------TRDQLTEEQIAEFKEAFSLFDKDGDGTITT
5303, 1631      KTMGWE-------------------------------------------------------TRDQLTEEQIAEFKEAFSLFDKDGDGTITT
5304, 1632      KTMGWEA------------------------------------------------------TRDQLTEEQIAEFKEAFSLFDKDGDGTITT
5305, 5300      KTMGWEAS-----------------------------------------------------TRDQLTEEQIAEFKEAFSLFDKDGDGTITT
5309            KTMGW-----------------------------------------------------LGGTRDQLTEEQIAEFKEAFSLFDKDGDGTITT
5339, 5340      KTMGWEA------------------------------------------------------TRDQLTEEQIAEFKEAFSLFDKDGDGTITT
5341, 5342      KTMGWEA------------------------------------------------------TRDQLTEEQIAEFKEAFSLFDKDGDGTITT
5306            KTMGWEASTERLYPEDGVLKGD---------------------------------------TRDQLTEEQIAEFKEAFSLFDKDGDGTITT
5307            KTMGWEASTERLYPEDGVLKGDIKMA-----------------------------------TRDQLTEEQIAEFKEAFSLFDKDGDGTITT
5308            KTMGWEASTERLYPEDGVLKGDIKMALRLKDGGRYLADFKTTYKAKKPVQMPGAYNVD---TRDQLTEEQIAEFKEAFSLFDKDGDGTITT
5310            KTMGWEASTERLYPEDGVLKG-------------------------------------LGGTRDQLTEEQIAEFKEAFSLFDKDGDGTITT
5311            KTMGWEASTERLYPEDGVLKGDIKM---------------------------------LGGTRDQLTEEQIAEFKEAFSLFDKDGDGTITT
5312            KTMGWEASTERLYPEDGVLKGDIKMALRLKDGGRYLADFKTTYKAKKPVQMPGAYNV-LGGTRDQLTEEQIAEFKEAFSLFDKDGDGTITT
Aa in mScarlet         147            162

K-GECO1         KELGTVMRSLGQNPTEAELQDMINEVDADGDGTFDFPEFLTMMARKMSYRVTEEEIREAFRVFDKDGNGYIGAAELRHVMTDLGEKLTDEE
R-GECO1         KELGTVMRSLGQNPTEAELQDMINEVDADGDGTFDFPEFLTMMARKMNDTDSEEEIREAFRVFDKDGNGYIGAAELRHVMTDLGEKLTDEE
CH-GECO2.1      KELGTVMRSLGQNPTEAELQDMINEVDADGDGTLDFPEFLTMMARKMNDSDSEEEIREAFRVFDKDGNGYIGAAELRHVMTNLGEKLTDEE
All others      KELGTVMRSLGQNPTEAELQDMINEVDADGDGTFDFPEFLTMMARKMNDTDSEEEIREAFRVFDKDGNGYIGAAELRHVMTDLGEKLTDEE

All             VDEMIRVADIDGDGQVNYEEFVQMMTAK*
```

**Not included in alignment**

5357 – as 5339, but with M164R
5358 – as 5339, but with M164K
5359 – as 5340, but with M164R
5360 – as 5340, but with M164K
5361 – as 5341, but with M164R
5362 – as 5341, but with M164K
5363 – as 5342, but with M164R
5364 – as 5342, but with M164K

**Legend**

amino acid – variations between variants
amino acid – relevant residues in mScarlet, numbers indicated
FMYGSR and FMYGSK - chromophore
PVV and TRD – linker 1 and 2 from R-GECO1
LES-LL and LGGTRD – linker 1 and 2 from CH-GECO2.1
KYNT and SNG – linker 1 and 2 from K-GECO1
S, M164R and M164K – point mutation
variant number - the first 10 designed variants

**Figure S1: Alignment of all mScarlet-I and mScarlet-A220 candidate sensors with three published red calcium sensors.** The variants are indicated by the numbers of our internal plasmid numbering system.

**Alignment all mScarlet3 and mScarlet-I3 variants**

```
K-GECO1         MASRMGSVKLIPSLTTVILVKSMLRKRSFGNPFKYNTETLYPADGGLEGACDMALKLVGGGHLNCSLETTYRSKKPATNLKMPGVYN
R-GECO1         -----MVDSSRRKWNKAGHAVRAIGRLSSPVV----SERMYPEDGALKSEIKKGLRLKDGGHYAAEVKTTYKAKKP---VQLPGAYI
CH-GECO2.1      -----MVDSSRRKWNKAGHAVRAIGRLSSLESLL-STERLYPEDGALKGEIKQRLKLKDGGHYAAEVKTTYKAKKP---VQLPDAYI
5309            -----MVDSSRRKWNKAGHAVRAIGRLSSLESLL-STERLYPEDGVLKGDIKMALRLKDGGRYLADFKTTYKAKKP---VQMPGAYN
5496,5497       -----MVDSSRRKWNKAGHAVRAIGRLSSLESLL-STERLYPEDVVLKGDIKMALRLKDGGRYLADFKTTYKAKKP---VQMPGAFN
5505,5506       -----MVDSSRRKWNKAGHAVRAIGRLSSLESLL-STERLYPEDVVLKGDIKMALRLKDGGRYLADFKTTYRAKKP---VQMPGAFN
5498            -----MVDSSRRKWNKAGHAVRAIGRLSSLESLL--TERLYPEDVVLKGDIKMALRLKDGGRYLADFKTTYKAKKP---VQMPGAFN
5499            -----MVDSSRRKWNKAGHAVRAIGRLSSLESLL--TERLYPEDVVLKGDIKMALRLKDGGRYLADFKTTYRAKKP---VQMPGAFN
5500,5507       -----MVDSSRRKWNKAGHAVRAIGRLSSPVV---STERLYPEDVVLKGDIKMALRLKDGGRYLADFKTTYKAKKP---VQMPGAFN
5502,5503       -----MVDSSRRKWNKAGHAVRAIGRLSSPVV---STERLYPEDVVLKGDIKMALRLKDGGRYLADFKTTYRAKKP---VQMPGAFN
5501            -----MVDSSRRKWNKAGHAVRAIGRLSSPVV----TERLYPEDVVLKGDIKMALRLKDGGRYLADFKTTYKAKKP---VQMPGAFN
5504            -----MVDSSRRKWNKAGHAVRAIGRLSSPVV----TERLYPEDVVLKGDIKMALRLKDGGRYLADFKTTYRAKKP---VQMPGAFN
5511            -----MVDSSRRKWNKAGHAVRAIGRLSSPVVWEASTERLYPEDVVLKGDIKMALRLKDGGRYLADFKTTYKAKKP---VQMPGAFN
5512            -----MVDSSRRKWNKAGHAVRAIGRLSSPVV--ASTERLYPEDVVLKGDIKMALRLKDGGRYLADFKTTYKAKKP---VQMPGAFN
5513,5514       -----MVDSSRRKWNKAGHAVRAIGRLSSLESLLASTERLYPEDVVLKGDIKMALRLKDGGRYLADFKTTYKAKKP---VQMPGAFN
5515            -----MVDSSRRKWNKAGHAVRAIGRLSSPVVWEASTERLYPEDVVLKGDIKMALRLKDGGRYLADFKTTYRAKKP---VQMPGAFN
Aa in mScarlet                                    147
                                                 L-S: mutagenesis in 5496, residues 29-30
                                                 EA:  mutagenesis in 5511, residues 29-30

K-GECO1         VDHRLERIKEADDETYVELHEVAVARYVGLGGG-----GGTGG-SVSE-------LIKENMPMKLYMEGTVNNHHFKCTSEGEGKPY
R-GECO1         VDIKLDIVSHNEDYTIVEQCERAEGRHST-GGMDELYKGGTGGSLVSKGEEDNMAIIKEFMRFKVHMEGSVNGHEFEIEGEGEGRPY
CH-GECO2.1      VDIKLDIVSHNEDYTIVEQYERAEGRHST-GGMDELYKGGTGGSMVSKGVEDNMAFIKEFMRFKVHMEGSVNGHEFEIEGEGEGRPY
5309            VDRKLDITSHNEDYTVVEQYERSEARHST-GGMDELYQGGSGG-MVSKGE----AVIKEFMRFKVHMEGSMNGHEFEIEGEGEGRPY
5496-5515       IDRKLDITSHNEDYTVVEQYERSVARHSTGGSGGSLYQGGSGG-MDS-TE----AVIKEFMRFKVHMEGSMNGHEFEIEGEGEGRPY
Aa in mScarlet   198                   220

K-GECO1         EGTQTMRIKVVEGGPLPFAFDILATSFMYGSRTFIKHPPGIPDFFKQSFPEGFTWERVTTYEDGGVLTATQDTSLQDGCLIYNVKVR
R-GECO1         EAFQTAKLKVTKGGPLPFAWDILSPQFMYGSKAYIKHPADIPDYFKLSFPEGFRWERVMNFEDGGIIHVNQDSSLQDGVFIYKVKLR
CH-GECO2.1      EGTQTAKLKVTKGGPLPFAWDILSPQFMYGSKAYIKHPADIPDYWKLSFPEGFKWERVMNFEDGGVVTVTQDSSLQDGEFIYKVKLR
5309            EGTQTAKLKVTKGGPLPFSWDILSPQFMYGSRAFTKHPADIPDYYKQSFPEGFKWERVMNFEDGGAVTVTQDTSLEDGTLIYKVKLR
5496,5497,5498,5500,5507,5501,5511,5512,5513
                EGTQTAKLKVTKGGPLPFSWDILSPQFMYGSRAFIKHPADIPDYWKQSFPEGFKWERVMIFEDGGTVSVTQDTSLEDGTLIYKVKLR
5505,5506,5499,5502,5503,5504,5514,5515
                EGTQTAKLRVTKGGPLPFSWDILSPQFMYGSRAFTKHPADIPDYWKQSFPEGFKWERVMNFEDGGAVSVAQDTSLEDGTLIYKVKLR
Aa in mScarlet                                    74

K-GECO1         GMNFPANGPVMQKKTLGWEA--SNGQLTEEQIAEFKEAFSLFDKDGDGTITTKELGTVMRSLGQNPTEAELQDMINEVDADGDGTFD
R-GECO1         GTNFPPDGPVMQKKTMGWEA--TRDQLTEEQIAEFKEAFSLFDKDGDGTITTKELGTVMRSLGQNPTEAELQDMINEVDADGDGTFD
CH-GECO2.1      GTNFPSNGPVMQKKTMGW-LGGTRDQLTEEQIAEFKEAFSLFDKGGAGTITTKELGTVMRSLGQNPTEAELQDMINEVDADGDGTLD
5309            GTNFPPDGPVMQKKTMGW-LGGTRDQLTEEQIAEFKEAFSLFDKDGDGTITTKELGTVMRSLGQNPTEAELQDMINEVDADGDGTFD
5496            GGNFPPDGPVMQKRTMGW-LGGTRDQLTEEQIAEFKEAFSLFDKDGDGTITTKELGTVMRSLGQNPTEAELQDMINEVDADGDGTFD
5497,5498       GGNFPPDGPVMQKRTMGWELGGTRDQLTEEQIAEFKEAFSLFDKDGDGTITTKELGTVMRSLGQNPTEAELQDMINEVDADGDGTFD
5505            GTNFPPDGPVMQKKTMGW-LGGTRDQLTEEQIAEFKEAFSLFDKDGDGTITTKELGTVMRSLGQNPTEAELQDMINEVDADGDGTFD
5506,5499       GTNFPPDGPVMQKKTMGWELGGTRDQLTEEQIAEFKEAFSLFDKDGDGTITTKELGTVMRSLGQNPTEAELQDMINEVDADGDGTFD
5500            GGNFPPDGPVMQKRTMGW----TRDQLTEEQIAEFKEAFSLFDKDGDGTITTKELGTVMRSLGQNPTEAELQDMINEVDADGDGTFD
5507,5501       GGNFPPDGPVMQKRTMGWE---TRDQLTEEQIAEFKEAFSLFDKDGDGTITTKELGTVMRSLGQNPTEAELQDMINEVDADGDGTFD
5502            GTNFPPDGPVMQKKTMGW----TRDQLTEEQIAEFKEAFSLFDKDGDGTITTKELGTVMRSLGQNPTEAELQDMINEVDADGDGTFD
5503,5504       GTNFPPDGPVMQKKTMGWE---TRDQLTEEQIAEFKEAFSLFDKDGDGTITTKELGTVMRSLGQNPTEAELQDMINEVDADGDGTFD
5511-5513       GGNFPPDGPVMQKRTMGW-LGGTRDQLTEEQIAEFKEAFSLFDKDGDGTITTKELGTVMRSLGQNPTEAELQDMINEVDADGDGTFD
5514,5515       GTNFPPDGPVMQKKTMGW-LGGTRDQLTEEQIAEFKEAFSLFDKDGDGTITTKELGTVMRSLGQNPTEAELQDMINEVDADGDGTFD
                                   LG: mutagenesis in 5496, residues 265-266
                                   LG: mutagenesis in 5511, residues 266-267

K-GECO1         FPEFLTMMARKMSYRVTEEEIREAFRVFDKDGNGYIGAAELRHVMTDLGEKLTDEEVDEMIRVADIDGDGQVNYEEFVQMMTAK*
R-GECO1         FPEFLTMMARKMNDTDSEEEIREAFRVFDKDGNGYIGAAELRHVMTDLGEKLTDEEVDEMIRVADIDGDGQVNYEEFVQMMTAK*
CH-GECO2.1      FPEFLTMMARKMNDSDSEEEIREAFRVFDKDGNGYIGAAELRHVMTNLGEKLTDEEVDEMIRVADIDGDGQVNYEEFVQMMTAK*
All others      FPEFLTMMARKMNDTDSEEEIREAFRVFDKDGNGYIGAAELRHVMTDLGEKLTDEEVDEMIRVADIDGDGQVNYEEFVQMMTAK*
```

**Not included in alignment**

All mutants of 5496 and 5511, but the positions of the mutations are indicated with boxes

**Legend**

- amino acid — variations between variants
- amino acid — relevant residues in mScarlet, numbers indicated
- FMYGSR and FMYGSK - chromophore
- PVV and TRD — linker 1 and 2 from R-GECO1
- LESLL and LGGTRD — linker 1 and 2 from CH-GECO2.1
- KYNT and SNG — linker 1 and 2 from K-GECO1
- variant number - the first 12 designed Scarlet3 and Scarlet-I3 variants

**Figure S2: Alignment of all mScarlet3 and mScarlet-I3 candidate sensors, with three published red calcium sensors and the best performing mScarlet-I sensor**. The variants are indicated by the numbers of our internal plasmid numbering system.

**Table S1: Fluorescence lifetime changes of mutagenesis variants of 5511 and 5496 in bacterial lysates.** Fluorescence lifetime was measured in bacterial lysate in the presence (0.1 mM $CaCl_2$) and absence (9.5 mM EDTA) of calcium, in *n* isolates at room temperature. The absolute phase ($\tau_\Phi$) and modulation lifetime ($\tau_M$) in the presence of calcium, and the difference in phase and modulation lifetime ($\Delta\tau_\Phi$ and $\Delta\tau_M$) between the two conditions (+$Ca^{2+}$ minus -$Ca^{2+}$) are reported, including standard deviation [sd] if applicable. Mutants are named after their ancestor and mutations, f.e. 5511mut C_I stems from variant 5511 and has mutations A30C and L266I, and 5496mut G_Y originates from variant 5496 with mutations S30G and L265Y. The original residues at the positions of mutations are indicated at the ancestors 5511 and 5496. Underlined variants are tested in HeLa cells. *These data are copied from **Table 1** for comparison with mutants.

| Internal number | Systematic name | $\tau_\varphi$ +$Ca^{2+}$ [sd] (ns) | $\tau_M$ +$Ca^{2+}$ [sd] (ns) | $\Delta\tau_\varphi$ [sd] (ns) | $\Delta\tau_M$ [sd] (ns) | n |
|---|---|---|---|---|---|---|
| *Mutagenesis of 5511, positions A30 and L266* | | | | | | |
| 5511 | <u>cp144ins144-RGL/CL-ScI3-GECO (A_L)*</u> | 3.20 [0.03] | 3.30 [0.05] | 0.35 [0.02] | 0.30 [0.01] | 12 |
| 4475 | <u>5511mut V_I</u> | 3.04 [0.02] | 2.62 [0.01] | 0.44 [0.01] | 0.37 [0.01] | 5 |
| | 5511mut V_V | 2.92 [0.02] | 2.59 [0.02] | 0.35 [0.01] | 0.28 [0.00] | 3 |
| 4476 | <u>5511mut C_I</u> | 3.14 [0.02] | 2.92 [0.02] | 0.23 [0.01] | 0.17 [0.01] | 4 |
| 4477 | <u>5511mut C_V</u> | 3.08 [0.01] | 2.92 [0.03] | 0.18 [0.03] | 0.13 [0.03] | 3 |
| | 5511mut C_T | 3.05 | 2.93 | 0.15 | 0.11 | 1 |
| | 5511mut L_V | 2.67 | 2.62 | 0.07 | 0.04 | 1 |
| | 5511mut W_T | 2.84 | 2.79 | 0.07 | 0.07 | 1 |
| *Mutagenesis of 5496, positions S30 and L265* | | | | | | |
| 5496 | <u>cp147del145-146_ScI3-CL-GECO (S_L)*</u> | 2.51 [0.06] | 2.72 [0.05] | -0.10 [0.01] | -0.03 [0.01] | 14 |
| | 5496mut S_Y | 2.44 [0.06] | 2.19 [0.04] | 0.24 [0.02] | 0.05 [0.00] | 2 |
| 4479 | <u>5496mut S_V</u> | 2.49 [0.02] | 2.71 [0.03] | -0.22 [0.01] | -0.14 [0.01] | 3 |
| 4478 | <u>5496mut S_I</u> | 2.53 [0.04] | 2.71 [0.05] | -0.19 [0.01] | -0.11 [0.01] | 5 |
| | 5496mut S_F | 2.61 [0.00] | 2.76 [0.00] | -0.14 [0.00] | -0.08 [0.00] | 2 |
| | 5496mut G_T | 2.37 [0.03] | 2.06 [0.03] | 0.30 [0.00] | 0.27 [0.00] | 5 |
| | 5496mut G_Q | 2.37 | 2.09 | 0.29 | 0.29 | 1 |
| | 5496mut G_E | 2.44 | 2.17 | 0.27 | 0.27 | 1 |
| | 5496mut G_H | 2.39 [0.02] | 2.15 [0.03] | 0.24 [0.01] | 0.23 [0.01] | 4 |
| 4481 | <u>5496mut G_F</u> | 2.38 [0.07] | 2.17 [0.06] | 0.22 [0.01] | 0.20 [0.01] | 3 |
| 4482 | <u>5496mut G_Y</u> | 2.39 [0.10] | 2.16 [0.10] | 0.22 [0.01] | 0.19 [0.01] | 2 |
| 4480 | <u>5496mut G_V</u> | 2.33 [0.03] | 2.13 [0.02] | 0.20 [0.00] | 0.20 [0.00] | 6 |
| | 5496mut G_A | 2.27 | 2.09 | 0.18 | 0.18 | 1 |
| | 5496mut G_C | 2.33 [0.06] | 2.15 [0.06] | 0.17 [0.01] | 0.17 [0.02] | 2 |
| 4483 | <u>5496mut G_L</u> | 2.41 | 2.24 | 0.16 | 0.16 | 1 |
| | 5496mut G_R | 2.14 | 2.06 | 0.09 | 0.08 | 1 |
| | 5496mut A_V | 2.19 | 2.04 | 0.15 | 0.18 | 1 |
| | 5496mut A_I | 2.07 | 2.00 | 0.07 | 0.12 | 1 |
| | 5496mut T_I | 2.95 | 2.91 | 0.03 | 0.03 | 1 |

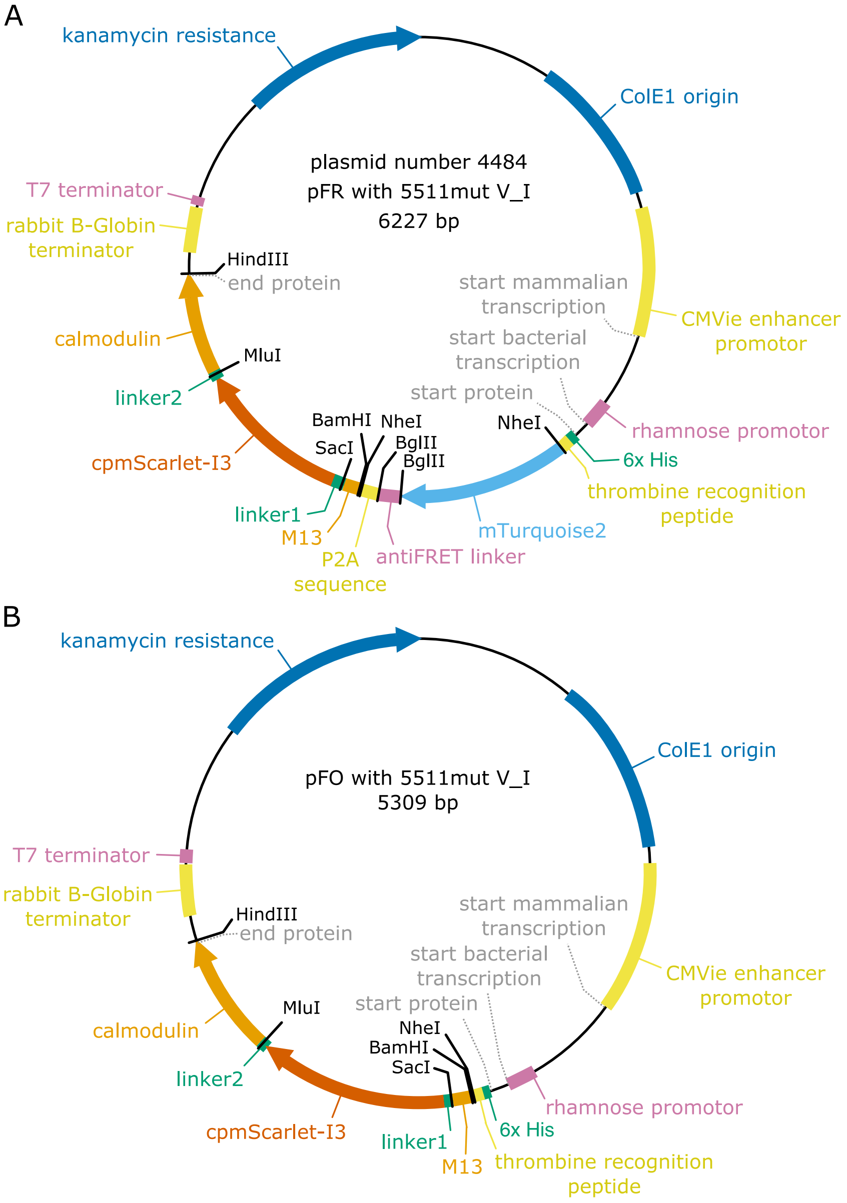


**Figure S3: Schematic representation of a pFR (Franka Ratio) and pFPO (Franka Protein Only) plasmid. A)** Expression of pFR in bacteria yields one protein containing: 6x His–thrombin recognition sequence–mTq2–antiFRET linker–P2A sequence–a red calcium sensor (M13 peptide, linker 1–circular permutated mScarlet–linker 2–calmodulin). In mammalian cells the protein in split in two at the P2A sequence. **B)** Expression of pFR in bacteria or mammalian cells yields one protein containing: 6x His–thrombin recognition sequence–a red calcium sensor (M13 peptide, linker 1–circular permutated mScarlet–linker 2–calmodulin).

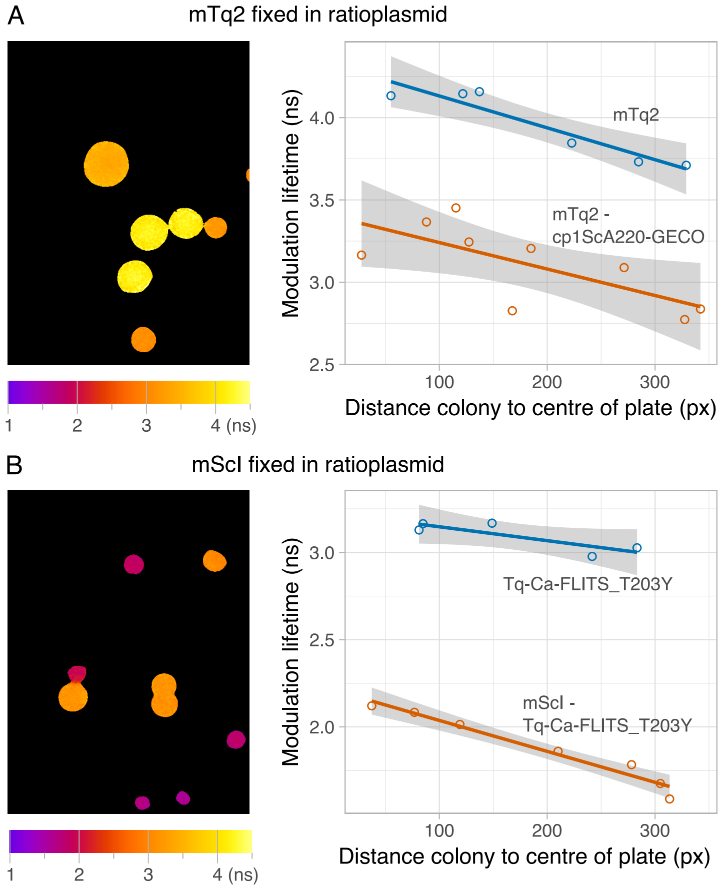


**Figure S4: Expression of the new ratioplasmids pFR in bacterial colonies. A)** A difference in modulation lifetime can be observed between colonies expressing mTq2 and colonies expressing mTq2 attached to a dummy red sensor via an anti-FRET linker. The left panel shows bacterial colonies with the false color indicating the modulation lifetime. The right panel shows the modulation lifetime of individual colonies (circles) plotted against the position of the colony on the agar plate. A linear model is fitted through the data (line) with a 95% confidence interval (gray area). A spread of the lifetime is observed, which is related to the distance from the colony to the middle of the agar plate. **B)** Same as **A** but comparing the modulation lifetime of Tq-Ca-FLITS_T203Y with and without mScI connected by an anti-FRET linker.

**Table S2: Influence of an RFP on the fluorescence lifetime of mTq2 and Tq-Ca-FLITS_T203Y in HeLa cells.** The fluorescence lifetimes were measured of the mTq2 or the Tq-Ca-FLITS_T203Y component of the indicated constructs, at 37 °C, before and after addition 5 µg/mL ionomycin and 5 mM $CaCl_2$ to the medium, and before and after photobleaching of the RFP to 22% of its original intensity. The average phase ($\Delta\tau_\phi$) and modulation lifetime changes ($\Delta\tau_M$) are calculated from the lifetime changes of individual cells ($n$). *The same bleaching pulse was applied as for the cells expressing an RFP.

| | +$Ca^{2+}$ minus -$Ca^{2+}$ | | | Bleaching RFP minus no bleaching | | |
|---|---|---|---|---|---|---|
| **Protein** | **$\Delta\tau_\varphi$ [sd] (ns)** | **$\Delta\tau_M$ [sd] (ns)** | **n** | **$\Delta\tau_\varphi$ [sd] (ns)** | **$\Delta\tau_M$ [sd] (ns)** | **n** |
| mTq2* | 0.05 [0.03] | 0.03 [0.02] | 11 | -0.01 [0.03] | -0.02 [0.03] | 30 |
| mTq2 and cp1mScA220-GECO | 0.04 [0.04] | 0.00 [0.05] | 6 | 0.01 [0.04] | 0.01 [0.03] | 11 |
| Tq-Ca-FLITS_T203Y* | -0.67 [0.05] | -0.78 [0.04] | 11 | -0.02 [0.06] | -0.03 [0.06] | 15 |
| Tq-Ca-FLITS_T203Y and mScI | -0.60 [0.06] | -0.72 [0.04] | 8 | -0.01 [0.03] | -0.01 [0.03] | 15 |

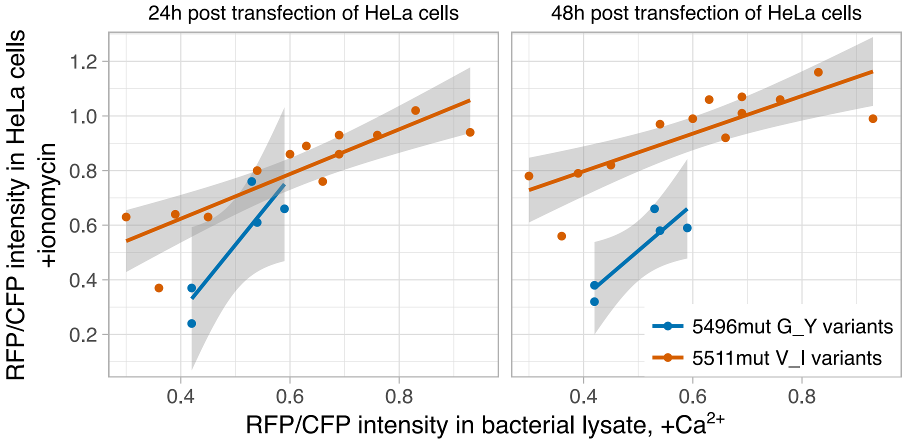


**Figure S5: Comparison of the intensity between bacterial lysate and HeLa cells of mutants of 5511mut V_I and 5496mut G_Y.** Bacteria and HeLa cells expressed simultaneously a red candidate sensor and mTq2 by using a pFR plasmid. Bacteria were lysed and the RFP/CFP intensity was measured in the presence (0.1 mM $CaCl_2$) of calcium. In HeLa cells the RFP/CFP intensity was measured after addition of of 5 µg/mL ionomycin and 5 mM $CaCl_2$. Each point indicates the average of the selected variant, which are taken from **Table S3** and **Figure 11**. A linear model is fitted through the data (line) with a 95% confidence interval (gray area).

**Table S3: Fluorescence changes of mutants of 5511mut V_I and 5496mut G_Y in bacterial lysates.** Bacteria expressed simultaneously a red candidate sensor and mTq2 coded on a pFR plasmid. Red and cyan fluorescence, and the fluorescence lifetime of the red candidate sensor was measured in bacterial lysate in the presence (0.1 mM $CaCl_2$) and absence (9.5 mM EDTA) of calcium, in *n* isolates at room temperature. Red fluorescence was divided over the cyan fluorescence (RFP/CFP). The average ratio in the calcium-bound state and the fold-change (+$Ca^{2+}$ / -$Ca^{2+}$) are indicated. For the lifetime, the absolute phase ($\tau_\Phi$) and modulation lifetime ($\tau_M$) in the presence of calcium, and the difference in phase and modulation lifetime ($\Delta\tau_\Phi$ and $\Delta\tau_M$) between the two conditions (+$Ca^{2+}$ minus -$Ca^{2+}$) are reported, including standard deviation [sd] if applicable. Mutants are named after their ancestor and mutations, f.e. 5511mut LV_IA stems from variant 5511mut V_I and has mutations E29L and G267A. The original residues at the positions of mutations are indicated at the ancestors 5511mut V_I and 5496mut G_Y. Underlined variants are tested in HeLa cells. *From these variants only the fluorescence lifetime could be measured, no mTq2 was expressed. **These two variants were the ancestors in the mutagenesis.

| Internal number | Systematic name | $\tau_\varphi$ +$Ca^{2+}$ [sd] (ns) | $\tau_M$ +$Ca^{2+}$ [sd] (ns) | $\Delta\tau_\varphi$ [sd] (ns) | $\Delta\tau_M$ [sd] (ns) | RFP/CFP +$Ca^{2+}$ | Fold change RFP/CFP | n |
|---|---|---|---|---|---|---|---|---|
| *Mutagenesis of 5511mut V_I, positions E29 and G267* | | | | | | | | |
| 5511 | cp144ins144-RGL/CL-ScI3-GECO (EA_LG)* | 3.25 [0.04] | 3.34 [0.03] | 0.35 [0.01] | 0.30 [0.01] | * | * | 3 |
| 4484† | 5511mut V_I (EV_IG)† | 3.13 [0.04] | 3.22 [0.03] | 0.44 [0.01] | 0.37 [0.01] | 0.36 | 1.3 | 3 |
| | 5511mut LV_IA | 3.07 [0.10] | 3.20 [0.06] | 0.61 [0.05] | 0.52 [0.03] | 0.30 | 1.3 | 2 |
| | 5511mut LV_IC | 3.07 | 3.19 | 0.61 | 0.51 | 0.60 | 1.4 | 1 |
| | 5511mut LV_IS | 3.13 [0.03] | 3.22 [0.02] | 0.60 [0.01] | 0.48 [0.00] | 0.54 | 1.4 | 3 |
| | 5511mut LV_IT | 3.10 | 3.22 | 0.58 | 0.49 | 0.63 | 1.4 | 1 |
| | 5511mut TV_IA | 3.01 | 3.14 | 0.60 | 0.54 | 0.36 | 1.4 | 1 |
| | 5511mut TV_IT T118I | 2.92 | 3.07 | 0.49 | 0.44 | 0.93 | 1.3 | 1 |
| | 5511mut SV_IG | 3.04 | 3.13 | 0.73 | 0.37 | 0.83 | 5.7 | 1 |
| | 5511mut SV_IA | 3.15 | 3.22 | 0.67 | 0.40 | 0.69 | 4.4 | 1 |
| | 5511mut SV_IT | 2.89 [0.02] | 3.02 [0.02] | 0.66 [0.01] | 0.38 [0.01] | 0.84 | 5.5 | 2 |
| | 5511mut SV_IS | 3.00 [0.05] | 3.10 [0.04] | 0.60 [0.01] | 0.32 [0.02] | 0.76 | 4.3 | 4 |
| | 5511mut SV_IH | 3.07 | 3.20 | ‡ | ‡ | 0.43 | 6.0 | 1 |
| | 5511mut SV_IF | 3.07 [0.02] | 3.19 [0.01] | ‡ | ‡ | 0.69 | 15.1 | 2 |
| | 5511mut SV_IW | 3.04 | 3.16 | ‡ | ‡ | 0.45 | 20.6 | 1 |
| | 5511mut SV_IP | 3.15 | 3.24 | ‡ | ‡ | 0.39 | 21.5 | 1 |
| | 5511mut CV_IH | 3.18 [0.07] | 3.27 [0.04] | 0.78 [0.02] | 0.29 [0.08] | 0.66 | 7.0 | 2 |
| | 5511mut CV_IS | 3.05 [0.05] | 3.15 [0.04] | 0.58 [0.03] | 0.31 [0.02] | 0.65 | 3.3 | 2 |
| | 5511mut CV_IN | 3.09 | 3.19 | 0.55 | 0.21 | 0.58 | 5.0 | 1 |
| | 5511mut AV_IF | 2.99 | 3.13 | 0.56 | 0.46 | 0.53 | 1.7 | 1 |
| | 5511mut AV_IW | 3.17 | 3.27 | 0.64 | 0.38 | 0.56 | 4.4 | 1 |
| | 5511mut VV_IY | 2.90 | 3.14 | 0.58 | 0.40 | 0.27 | 1.5 | 1 |
| *Mutagenesis of 5496mut G_Y, positions L29 and G266* | | | | | | | | |
| 5496 | cp147del145-146_ScI3-CL-GECO (LS_LG)* | 2.52 [0.01] | 2.73 [0.00] | -0.09 [0.02] | -0.04 [0.01] | * | * | 3 |
| 4491† | 5496mut G_Y (LG_YG) † | 2.39 [0.01] | 2.58 [0.00] | 0.26 [0.02] | 0.19 [0.01] | 0.59 | 1.6 | 3 |
| | 5496mut IG_YD | 2.52 | 2.69 | 0.51 | 0.40 | 0.18 | 1.1 | 1 |
| | 5496mut IG_YM | 2.49 | 2.69 | 0.48 | 0.39 | 0.51 | 1.5 | 1 |
| | 5496mut IG_YL | 2.50 | 2.69 | 0.45 | 0.39 | 0.41 | 1.8 | 1 |
| | 5496mut IG_YP | 2.56 [0.05] | 2.75 [0.03] | 0.45 [0.01] | 0.38 [0.01] | 0.46 | 1.2 | 3 |
| | 5496mut LG_YY | 0.99 | 1.75 | 0.39 | 0.55 | 0.22 | 1.3 | 1 |
| | 5496mut LG_YL | 2.45 [0.01] | 2.67 [0.01] | 0.49 [0.01] | 0.42 [0.01] | 0.42 | 1.7 | 3 |
| | 5496mut LG_YP | 2.45 [0.03] | 2.68 [0.02] | 0.48 [0.02] | 0.42 [0.02] | 0.54 | 1.3 | 8 |
| | 5496mut VG_YM | 2.48 [0.01] | 2.67 [0.00] | 0.51 [0.00] | 0.41 [0.00] | 0.42 | 1.7 | 2 |
| | 5496mut VG_YP | 2.54 [0.02] | 2.73 [0.01] | 0.45 [0.01] | 0.38 [0.02] | 0.45 | 1.2 | 2 |
| | 5496mut FG_YA | 2.42 | 2.66 | 0.53 | 0.34 | 0.53 | 1.5 | 1 |
| | 5496mut FG_YL | 2.47 | 2.67 | 0.44 | 0.32 | 0.51 | 2.0 | 1 |
| | 5496mut MG_YP | 2.36 | 2.58 | 0.45 | 0.37 | 0.36 | 1.4 | 1 |

**Table S4: Fluorescence lifetime changes of RCaMP1h and jRCaMP1b in bacterial lysate buffer.** Fluorescence lifetime was measured in the presence (0.1 mM $CaCl_2$) and absence (9.5 mM EDTA) of calcium, in duplo at room temperature. Calcium sensors were obtained by crude extraction using the bacterial lysis protocol[23], or by protein isolation using a Ni-NTA column in combination with a His-tag. Proteins were diluted in Tris-HCl pH 8.0 buffer with different concentrations of DOC. 0.8% DOC was used for measurements of the bacterial screening of 5511mut V_I and 5496mut G_Y, 2% DOC was used for all other measurements in bacterial lysate.

| | | | Phase lifetime (ns) [sd] | | | Modulation lifetime (ns) [sd] | | |
|---|---|---|---|---|---|---|---|---|
| **Sensor** | **Isolation type** | **DOC (%)** | **+$Ca^{2+}$** | **-$Ca^{2+}$** | $\Delta\tau_{\varphi}$ | **+$Ca^{2+}$** | **-$Ca^{2+}$** | $\Delta\tau_{M}$ |
| jRCaMP1b | bacteria lysate, pFR | 0.8% | 2.48 [0.00] | 2.75 [0.00] | -0.27 [0.00] | 2.98 [0.00] | 3.10 [0.00] | -0.12 [0.00] |
| RCaMP1h | bacteria lysate, pFR | 0.8% | 2.26 [0.03] | 1.48 [0.00] | 0.78 [0.04] | 2.86 [0.02] | 2.14 [0.01] | 0.72 [0.03] |
| RCaMP1h | protein isolate | 0% | 3.11 [0.00] | 1.15 [0.00] | 1.96 [0.00] | 3.22 [0.00] | 1.61 [0.00] | 1.61 [0.01] |
| RCaMP1h | protein isolate | 0.8% | 2.04 [0.00] | 1.59 [0.01] | 0.45 [0.01] | 2.70 [0.00] | 2.25 [0.01] | 0.45 [0.01] |
| RCaMP1h | protein isolate | 2% | 1.94 [0.00] | 1.86 [0.00] | 0.08 [0.00] | 2.61 [0.00] | 2.50 [0.00] | 0.11 [0.00] |

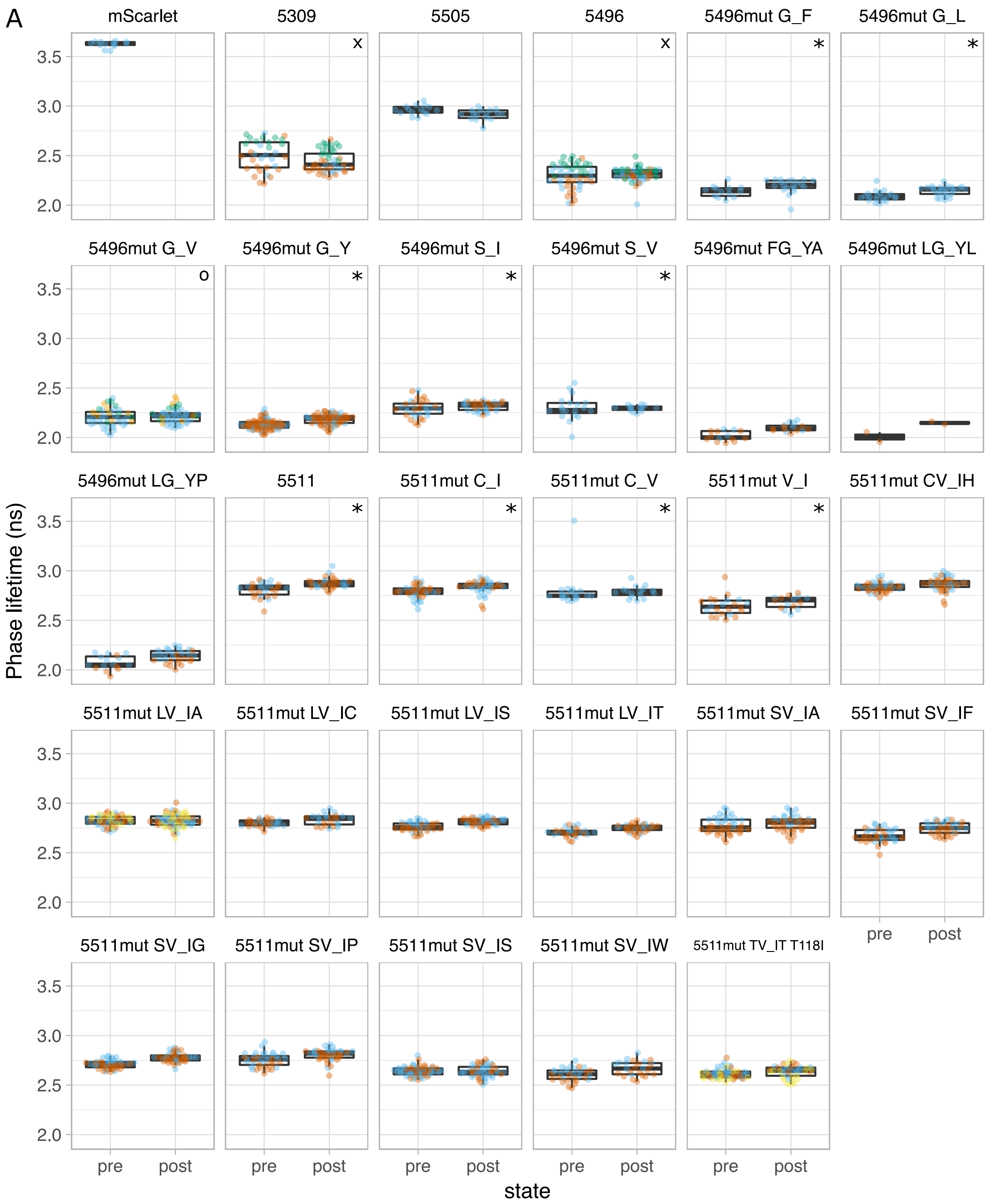


**Figure S6: Fluorescence lifetime measurements of HeLa cells expressing selected candidate sensors.** The fluorescence phase (**A**) and modulation (**B**) lifetime were measured at 37 °C, before (pre) and after (post) addition of 5 µg/mL ionomycin and 5 mM $CaCl_2$. Each point indicates a single cell. Different colors indicate different microscopy glasses. Variants 5309, 5496 and 5496mut G_V were measured on two different days. Boxplots indicate the median (line), upper and lower quartile (box) and 1.5× interquartile range measured from the lower and upper quartiles (whiskers). *: Pseudo random phase recording order was off for all measurements, O: Pseudo random phase recording order was off for measurements in blue, x: Pseudo random phase recording order was off for measurements in red and blue

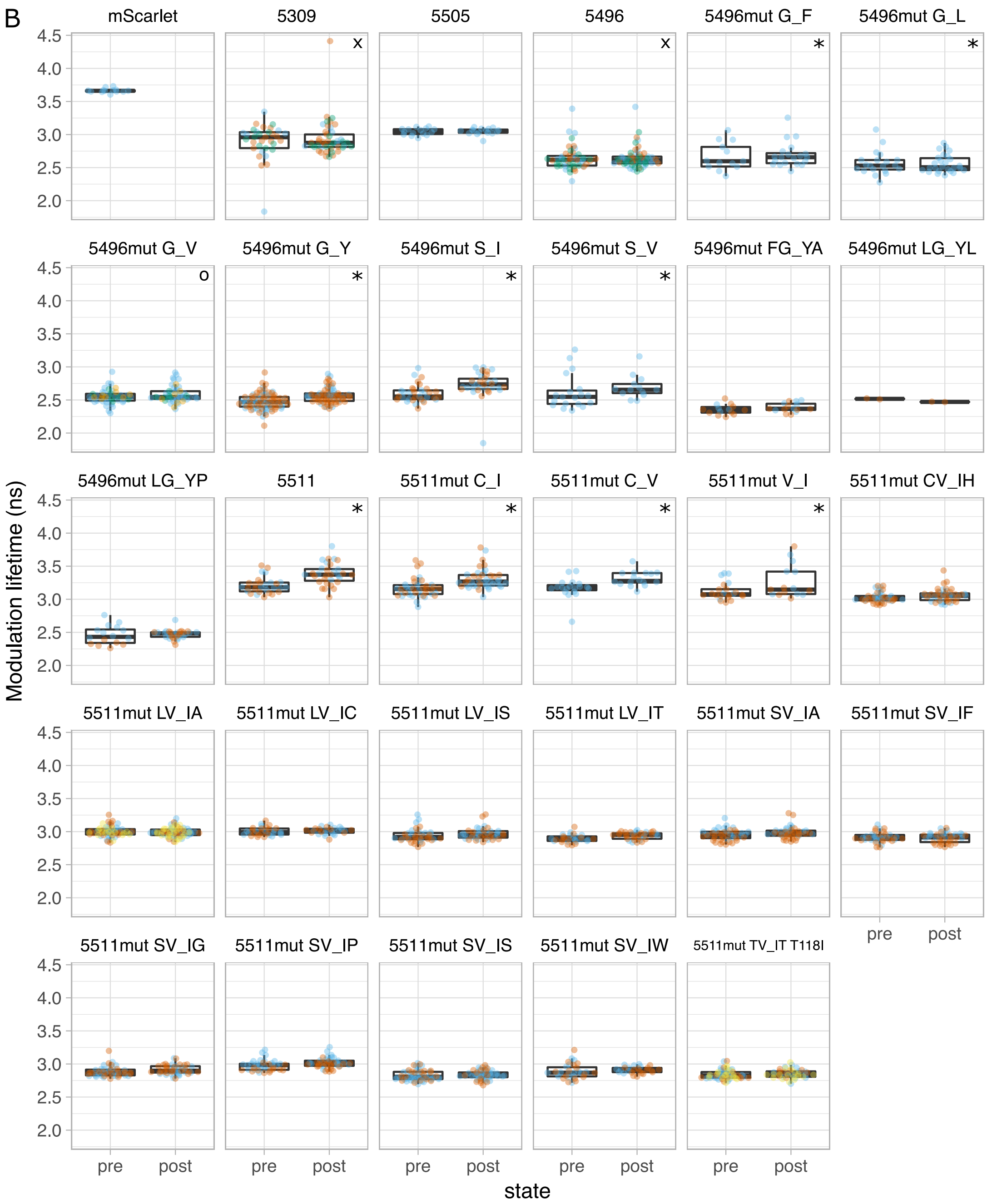


**Figure S6:** (continued)

**Table S5: Primers used in this study.** All primers were ordered from Integrated DNA Technologies.

| No. | Name | Sequence |
|---|---|---|
| 1 | FW cp144 mSIRG | GCTGAGCTCACCCGTGGTTTGGGAAGCATCCACCGAG |
| 2 | RV cp144 mSIRG | GTCACGCGTGCCCATTGTCTTCTTCTG |
| 3 | FW cp145 mSIRG | GCTGAGCTCACCCGTGGTTGAAGCATCCACCGAGCG |
| 4 | RV cp145 mSIRG | GTCACGCGTCCAGCCCATTGTCTTCTTC |
| 5 | FW cp146 mSIRG | GCTGAGCTCACCCGTGGTTGCATCCACCGAGCGG |
| 6 | RV cp146 mSIRG | GTCACGCGTTTCCCAGCCCATTGTCTTC |
| 7 | FW cp147 mSIRG | GCTGAGCTCACCCGTGGTTTCCACCGAGCGGTTGTAC |
| 8 | RV cp147 mSIRG | GTCACGCGTTGCTTCCCAGCCCATTG |
| 9 | FW cp148 mSIRG | GCTGAGCTCACCCGTGGTTACCGAGCGGTTGTACCC |
| 10 | RV cp148 mSIRG | GTCACGCGTGGATGCTTCCCAGCCC |
| 11 | FW cpT148S mSRG | GCTGAGCTCACCCGTGGTTAGCGAGCGGTTGTACCC |
| 12 | FW cp162 mSRG | GCTGAGCTCACCCGTGGTTATTAAGATGGCCCTGCGC |
| 13 | RV cp162 mSRG | GTCACGCGTGTCGCCCTTCAGCACGC |
| 14 | FW cp166 mSRG | GCTGAGCTCACCCGTGGTTCTGCGCCTGAAGGACGG |
| 15 | RV cp166 mSRG | GTCACGCGTGGCCATCTTAATGTCGCC |
| 16 | FW cp198 mSIRG | GCTGAGCTCACCCGTGGTTCGCAAGTTGGACATCACC |
| 17 | RV cp198 mSIRG | GTCACGCGTGTCGACGTTGTAGGCGC |
| 18 | FW_GECO-SCL1 | GCTGAGCTCACTTGAGTCGCTGCTGTCCACCGAGCGGTTGTAC |
| 19 | RV_GECO-SCL1 | GTCACGCGTACCGCCCAGCCAGCCCATTGTCTTCTTC |
| 20 | FW_GECO-SCL2 | GCTGAGCTCACTTGAGTCGCTGCTGATTAAGATGGCCCTGCGC |
| 21 | RV_GECO-SCL2 | GTCACGCGTACCGCCCAGGCCCTTCAGCACGCCG |
| 22 | FW_GECO-SCL3 | GCTGAGCTCACTTGAGTCGCTGCTGCTGCGCCTGAAGGACGG |
| 23 | RV_GECO-SCL3 | GTCACGCGTACCGCCCAGCATCTTAATGTCGCCCTTC |
| 24 | FW_GECO-SCL4 | GCTGAGCTCACTTGAGTCGCTGCTGCGCAAGTTGGACATCACC |
| 25 | RV_GECO-SCL4 | GTCACGCGTACCGCCCAGGACGTTGTAGGCGCCGG |
| 26 | FW M164K/R mSc | GGCGACATTAAGARGGCCCTGCGCCTG |
| 27 | RV M164K/R mSc | CAGGCGCAGGGCCYTCTTAATGTCGCC |
| 28 | FW cp148 mSKL IVA | GCTGAGCTCAAAGTATAATACCGAGCGGTTGTACCCC |
| 29 | RV cp148 mSKL IVA | AGTCAATTGGCCATTACTTGCTTCCCAGCCCATTG |
| 30 | FW backbone GKL IVA | AGTAATGGCCAATTGACTGAAGAGCAGATC |
| 31 | RV backbone GKL IVA | ATTATACTTTGAGCTCAGCCGACCTATAG |
| 32 | FW cp1_from_td | GCTGAGCTCACCCGTGGTTATGGTGAGCAAGGGCGAG |
| 33 | RV cp1_from_td | GTCACGCGTCTGGTACAGCTCGTCCATG |
| 34 | RV_cp146-Sc3CL | GTCACGCGTACCGCCCAGTTCCCAGCCCATTGTCTTC |
| 35 | FW_cp148-Sc3CL | GCTGAGCTCACTTGAGTCGCTGCTGACCGAGCGGTTGTACCC |
| 36 | RV cp145 mScI3RG | GTCACGCGTCCAGCCCATTGTCCTCTTC |
| 37 | RV cp146 mScI3RG | GTCACGCGTTTCCCAGCCCATTGTCCTC |
| 38 | RV_cp145-ScI3CL | GTCACGCGTACCGCCCAGCCAGCCCATTGTCCTCTTC |
| 39 | RV_cp146-ScI3CL | GTCACGCGTACCGCCCAGTTCCCAGCCCATTGTCCTC |
| 40 | FW_cp146-Sc3CL | GCTGAGCTCACTTGAGTCGCTGCTGGCATCCACCGAGCGG |
| 41 | FW_WEX_cpSc(I)3 | GCTGAGCTCACCCGTGGTTTGGGAANNKTCCACCGAGCGGTTGTAC |
| 42 | RV_cpScI3_XGG | GTCACGCGTACCGCCMNNCCAGCCCATTGTCCTCTTC |
| 43 | FW_LLX_cpSc(I)3 | GCTGAGCTCACTTGAGTCGCTGCTGNNKACCGAGCGGTTGTACCC |
| 44 | FW_198I_ScI3 | GCCTTCAACATCGACATCAAGTTGGACATCACC |
| 45 | RV_198I_ScI3 | GGTGATGTCCAACTTGATGTCGATGTTGAAGGC |
| 46 | FW_LXG_cpSc(I)3 | GCTGAGCTCACTTGAGTCGCTGNNKGGGACCGAGCGGTTGTAC |
| 47 | RV_cpScI3_YXG | GTCACGCGTACCMNNATACCAGCCCATTGTCC |
| 48 | FW_WXV_cpSc(I)3 | GCTGAGCTCACCCGTGGTTTGGNNKGTGTCCACCGAGCGGTTG |
| 49 | RV_cpScI3_IXG | GTCACGCGTACCMNNAATCCAGCCCATTGTCCTC |
| 50 | FW-del_SacI_in-FP-pDress | CGGCATGGACGAGCTGTACAAGTCGACCAG |
| 51 | RV-del_SacI_in-FP-pDress | CTGGTCGACTTGTACAGCTCGTCCATGCCG |
| 52 | FW_FP-lessFRET-P2A | GTCGCGAATTCAGGCGC |
| 53 | RV_FP-lessFRET-P2A | TTTGGATCCAGCGCTAGCGAAGGTCC |
| 54 | FW_FLITS-backbone_notags | TAAGGATCCCGCCACAATGGTC |
| 55 | RV_FLITS_backbone_notags | GCGCCTGAATTCGCGAC |
| 56 | FW_KGECO_to_ratio | GCTGGATCCCGCCACAATGGGCAGCGTGAAGC |
| 57 | RV_RCaMPs_to_ratio | CACAAGCTTCTACTTCGCTGTCATCATTTG |
| 58 | FW_RCaMPs_to_ratio | GCTGGATCCCGCCACAATGGTCGACTCATCACG |
| 59 | FW_FP_to_pFR | GCTGGATCCCGCCACCATGGTGAGC |
| 60 | RV_FP_to_pFR | CACAAGCTTTTACTTGTACAGCTCGTCC |
| seq1 | FW seq_FP_in_GECO | GTGGACAGCAAATGGGTCG |
| seq2 | RV_seq_FP_in_GECO | CCCAGAGACCGCATCACC |
| seq3 | #09: FruitFPout | CATCACCTCCCACAACG |
| seq4 | #12: 3'-pBABEseq | ACCCTAACTGACACACATTCC |
| seq5 | RV cp1_from_td | GTCACGCGTCTGGTACAGCTCGTCCATG |
| seq6 | FW_pFH_on_pDXterm | GCGAAGTAGAAGCTTGTGATTAACCTCAGGTGCAG |
| seq7 | RV925 | CGATCTCAGTGGTATTTGTGAG |
| seq8 | FW45 | GCTTTTTAGACTGGTCGTAGGGA |
| seq9 | FW cp149mTQ2RG | GCTGAGCTCACCCGTGGTTAACGTCTATATCACCGCC |

**Table S6: Overview of all variants constructed in this study.** The plasmid numbers stem from our internal plasmid numbering system.

| Plasmid number | Name | Fluorescent protein | linker type | FW primer | RV primer | Notes |
|---|---|---|---|---|---|---|
| *First mScarlet-I and mScarlet-A220 variants (10 in total)* | | | | | | |
| 1629 | pFHL-torPE-cp144mScarlet-I-GECO1 | mScarlet-I | R-GECO | 1 | 2 | |
| 1630 | pFHL-torPE-cp145mScarlet-I-GECO1 | mScarlet-I | R-GECO | 3 | 4 | |
| 1631 | pFHL-torPE-cp146mScarlet-I-GECO1 | mScarlet-I | R-GECO | 5 | 6 | |
| 1632 | pFHL-torPE-cp147mScarlet-I-GECO1 | mScarlet-I | R-GECO | 7 | 8 | |
| 5300 | pFHL-torPE-cp148mScarlet-I-GECO1 | mScarlet-I | R-GECO | 9 | 10 | |
| 5301 | pFHL-torPE-cp144mScarlet-A220-GECO1 | mScarlet-A220 | R-GECO | 1 | 2 | |
| 5302 | pFHL-torPE-cp145mScarlet-A220-GECO1 | mScarlet-A220 | R-GECO | 3 | 4 | |
| 5303 | pFHL-torPE-cp146mScarlet-A220-GECO1 | mScarlet-A220 | R-GECO | 5 | 6 | |
| 5304 | pFHL-torPE-cp147mScarlet-A220-GECO1 | mScarlet-A220 | R-GECO | 7 | 8 | |
| 5305 | pFHL-torPE-cp148mScarlet-A220-GECO1 | mScarlet-A220 | R-GECO | 9 | 10 | |
| *Control: dummy sensor* | | | | | | |
| 5313 | pFHL-torPE-cp1mScarlet-A220-GECO1 | mScarlet-A220 | R-GECO | 32 | 33 | pFR version is #5521 |
| *Additional mScarlet-I and mScarlet-A220 variants: deletions, mutations, other positions of cp (21 in total)* | | | | | | |
| 5306 | pFHL-torPE-cp162mScarlet-A220-GECO1 | mScarlet-A220 | R-GECO | 12 | 13 | |
| 5307 | pFHL-torPE-cp166mScarlet-A220-GECO1 | mScarlet-A220 | R-GECO | 14 | 15 | |
| 5308 | pFHL-torPE-cp198mScarlet-A220-GECO1 | mScarlet-A220 | R-GECO | 16 | 17 | |
| 5309 | pFHL-torPE-cp147del145-146-CL-mScarlet-A220-GECO | mScarlet-A220 | CH-GECO | 18 | 19 | pFR version is #4474 |
| 5310 | pFHL-torPE-cp162del161-CL-mScarlet-A220-GECO | mScarlet-A220 | CH-GECO | 20 | 21 | |
| 5311 | pFHL-torPE-cp166del165-CL-mScarlet-A220-GECO | mScarlet-A220 | CH-GECO | 22 | 23 | |
| 5312 | pFHL-torPE-cp198del197-CL-mScarlet-A220-GECO | mScarlet-A220 | CH-GECO | 24 | 25 | |
| 5339 | pFHL-torPE-cp148del147mScarlet-I GECO | mScarlet-I | R-GECO | 9 | 8 | |
| 5340 | pFHL-torPE-cp148del147mScarlet-A220 GECO | mScarlet-A220 | R-GECO | 9 | 8 | |
| 5341 | pFHL-torPE-cpT148Sdel147mScarlet-I GECO | mScarlet-I | R-GECO | 11 | 8 | #5339 with T148S |
| 5342 | pFHL-torPE-cpT148Sdel147mScarlet-A220 GECO | mScarlet-A220 | R-GECO | 11 | 8 | #5340 with T148S |
| 5343 | pFHL-torPE-cp148del147-KL-mScarlet-I-GECO | mScarlet-I | K-GECO | 28, 30 | 29, 31 | constructed by IVA |
| 5344 | pFHL-torPE-cp148del147-KL-mScarlet-A220-GECO | mScarlet-A220 | K-GECO | 28, 30 | 29, 31 | constructed by IVA |
| 5357 | pFHL-torPE cp148del147 M164R mScarlet-I GECO | mScarlet-I | R-GECO | 26 | 27 | #5339 with M164R |
| 5358 | pFHL-torPE cp148del147 M164K mScarlet-I GECO | mScarlet-I | R-GECO | 26 | 27 | #5339 with M164K |
| 5359 | pFHL-torPE cp148del147 M164R mScarlet-A220 GECO | mScarlet-A220 | R-GECO | 26 | 27 | #5340 with M164R |
| 5360 | pFHL-torPE cp148del147 M164K mScarlet-A220 GECO | mScarlet-A220 | R-GECO | 26 | 27 | #5340 with M164K |
| 5361 | pFHL-torPE-cpT148Sdel147 M164K mScarlet-I GECO | mScarlet-I | R-GECO | 26 | 27 | #5339 with T148S and M164K |
| 5362 | pFHL-torPE-cpT148Sdel147 M164R mScarlet-I GECO | mScarlet-I | R-GECO | 26 | 27 | #5339 with T148S and M164R |
| 5363 | pFHL-torPE-cpT148Sdel147 M164K mScarlet-A220 GECO | mScarlet-A220 | R-GECO | 26 | 27 | #5340 with T148S and M164K |
| 5364 | pFHL-torPE-cpT148Sdel147 M164R mScarlet-A220 GECO | mScarlet-A220 | R-GECO | 26 | 27 | #5340 with T148S and M164R |
| *mScarlet3 and mScarlet-I3 variants, based on variant 5309 (12 in total)* | | | | | | |
| 5505 | pFHL-TorPE-cp147del145-146_Sc3-CL-GECO | mScarlet3 | CH-GECO | 18 | 19 | |
| 5506 | pFHL-TorPE-cp147del146_Sc3-CL_GECO | mScarlet3 | CH-GECO | 18 | 34 | |
| 5499 | pFHL-TorPE-cp148del146-147_Sc3-CL-GECO | mScarlet3 | CH-GECO | 35 | 34 | |
| 5500 | pFHL-TorPE-cp147del145-146_ScI3-GECO | mScarlet-I3 | R-GECO | 7 | 36 | |
| 5507 | pFHL-TorPE-cp147del146_ScI3-GECO | mScarlet-I3 | R-GECO | 7 | 37 | |
| 5501 | pFHL-TorPE-cp148del146-147_ScI3-GECO | mScarlet-I3 | R-GECO | 9 | 37 | |
| 5502 | pFHL-TorPE-cp147del145-146_Sc3-GECO | mScarlet3 | R-GECO | 7 | 4 | |
| 5503 | pFHL-TorPE-cp147del146_Sc3-GECO | mScarlet3 | R-GECO | 7 | 6 | |
| 5504 | pFHL-TorPE-cp148del146-147_Sc3-GECO | mScarlet3 | R-GECO | 9 | 6 | |
| 5496 | pFHL-TorPE-cp147del145-146_ScI3-CL-GECO | mScarlet-I3 | CH-GECO | 18 | 38 | pFR version is #4473 |
| 5497 | pFHL-TorPE-cp147del146_ScI3-CL_GECO | mScarlet-I3 | CH-GECO | 18 | 39 | |
| 5498 | pFHL-TorPE-cp148del146-147_ScI3-CL-GECO | mScarlet-I3 | CH-GECO | 35 | 39 | |
| *mScarlet3 and mScarlet-I3 variants, screening of mixed PCR (100 new possibilities in total)* | | | | | | |
| 5511 | pFHL-torPE-cp144ins144-RGL/CL-ScI3-GECO | mScarlet-I3 | R/CH-GECO | | | pFR version is #4472 |
| 5512 | pFHL-torPE-cp146del145-RGL/CL-ScI3-GECO | mScarlet-I3 | R/CH-GECO | | | |
| *mScarlet3 and mScarlet-I3 variants, additional design (3 in total)* | | | | | | |
| 5513 | pFHL-torPE-cp146del145-CL-ScI3-GECO | mScarlet-I3 | CH-GECO | 40 | 38 | |
| 5514 | pFHL-torPE-cp146del145-CL-Sc3-GECO | mScarlet3 | CH-GECO | 40 | 19 | |
| 5515 | pFHL-torPE-cp144ins144-RGL/CL-Sc3-GECO | mScarlet3 | R/CH-GECO | 1 | 19 | |
| Variants based on 5511 and 5496 | | | | | | |
| 4475 | pFHL-TorPE-5511mutVI | mScarlet-I3 | R/CH-GECO | 41 | 42 | pFR version is #4484 |
| 4476 | pFHL-TorPE-5511mutCI | mScarlet-I3 | R/CH-GECO | 41 | 42 | pFR version is #4485 |
| 4477 | pFHL-TorPE-5511mutCV | mScarlet-I3 | R/CH-GECO | 41 | 42 | pFR version is #4486 |
| 4478 | pFHL-TorPE-5496mutSI | mScarlet-I3 | CH-GECO | 43 | 42 | pFR version is #4487 |
| 4479 | pFHL-TorPE-5496mutSV | mScarlet-I3 | CH-GECO | 43 | 42 | pFR version is #4488 |
| 4480 | pFHL-TorPE-5496mutGV | mScarlet-I3 | CH-GECO | 43 | 42 | pFR version is #4489 |
| 4481 | pFHL-TorPE-5496mutGF | mScarlet-I3 | CH-GECO | 43 | 42 | pFR version is #4490 |
| 4482 | pFHL-TorPE-5496mutGY | mScarlet-I3 | CH-GECO | 43 | 42 | pFR version is #4491 |
| 4483 | pFHL-TorPE-5496mutGL | mScarlet-I3 | CH-GECO | 43 | 42 | pFR version is #4492 |
| 4496 | pFR-mTq2-5496-GY-198I | mScarlet-I3 | CH-GECO | 44 | 45 | #4482/#4491 with R198I |
| 4497 | pFR-mTq2-5496-SV-198I | mScarlet-I3 | CH-GECO | 44 | 45 | #4479/#4488 with R198I |
| 4498 | pFR-mTq2-5511-CI-198I | mScarlet-I3 | R/CH-GECO | 44 | 45 | #4476/#4485 with R198I |
| 4499 | pFR-mTq2-5511-VI-198I | mScarlet-I3 | R/CH-GECO | 44 | 45 | #4475/#4484 with R198I |
| xxxxx | pFR-mTq2-5496mut XG_YX | mScarlet-I3 | CH-GECO | 46 | 47 | All variants of the mutagenesis of 5496mut G_Y |
| xxxxx | pFR-mTq2-5511mut XV_IX | mScarlet-I3 | R/CH-GECO | 48 | 49 | All variants of the mutagenesis of 5511mut V_I |
